# Hepatic stellate cell FXR signaling regulates context-dependent functions in liver homeostasis and fibrosis

**DOI:** 10.64898/2026.08.29.747537

**Authors:** Manjula Vinod, Francesco Paolo Zummo, Céline Gheeraert, Zouriatou Gouda, Sandra Courquet, Emilie Dorchies, Léna Thuret, Marie Lapage, Loic Guille, Marie Bobowski-Gérard, Charlène Pourpe, Victor Launay, Mehdi Derhoudi, Amélie Bonnefond, Delphine Eberlé, Joel T. Haas, Julie Dubois-Chevalier, Jérôme Eeckhoute, Sophie Lestavel, Bart Staels, Philippe Lefebvre, Alexandre Berthier

## Abstract

Nuclear bile acid (BA) signaling plays a central role in liver homeostasis and represents a major therapeutic axis in fibrotic liver diseases. The farnesoid X receptor (FXR), a master nuclear effector of BA signaling, is expressed in several liver-resident cell types, suggesting that it may regulate distinct biological programs beyond the hepatocyte (HC) compartment. Using complementary pharmacological, genetic, and computational approaches across in vitro, ex vivo, and in vivo models of mouse and human origin, we investigated the role of hepatic stellate cell (HSC) FXR (FXR^HSC^) in both unchallenged and injured livers, which has remained controversial. FXRα is robustly expressed in both HCs and HSCs with distinct isoform distributions, and these isoforms exhibited differential capacities to activate gene expression in an HSC context. We found that the potent selective FXR agonist tropifexor triggers a transcriptional program reminiscent of that observed after partial hepatectomy and associated with HC proliferation. This cell cycle-related response was also observed in HSCs and did not require intestinal FXR expression. An HSC-specific response to tropifexor was observed for several genes, including members of the glutathione-S-transferase (GST) family or *Scube1*. FXR^HSC^ was sufficient to observe the anti-fibrotic effects of tropifexor in precision-cut liver slices, an ex-vivo model of fibrosis. Finally, we identified the regulation of the chemerin-encoding gene *Rarres2* as a relevant example of FXR^HSC^-dependent control of hepatic intercellular communication. Together, these findings identify FXR^HSC^ as an important contributor to hepatic adaptation and therapeutic response to BA analogs and confirmed HSCs as a significant site of nuclear bile acid signaling in liver biology.

## INTRODUCTION

Nuclear bile acid (BA) signaling relies on a sophisticated regulatory network that orchestrates various metabolic processes critical for maintaining physiological homeostasis. At the core of this intricate system lies the farnesoid X receptor (FXR), a nuclear receptor pivotal in BA metabolism and beyond [1, 2]. FXR expression distribution is well defined and shows prominent expression in the liver and ileum, and significant amounts are also detected in the kidney. Loss-of-function approaches have also suggested a role for FXR-driven signaling in the pancreas, adrenal glands and adipose tissue [3].

Studies have primarily investigated the metabolic functions of liver FXR, with an initial emphasis on the molecular dissection of the regulation of the metabolism of cholesterol derivatives, BAs [4]. As a nuclear BA sensor, FXR (encoded by the *Nr1h4* gene) limits BA accumulation within hepatocytes (HCs) by promoting BAs export and repressing their synthesis. This dual action mitigates BA accumulation within HCs, thus safeguarding against BA-induced hepatocellular injury and inflammation, thereby maintaining hepatic integrity through a mechanism implying intestinal FXR [3]. Furthermore, liver FXR regulates glucose, amino acid, lipid metabolism, and autophagy, thereby establishing FXR as a fundamental transcription factor in maintaining overall hepatic energy and metabolic balance [2, 5]. The vast majority of FXR activities have thus been ascribed to hepatocyte FXR (FXR^HC^), and initial studies were mostly conducted in HC-like cellular models or in whole body or liver-specific *Nr1h4*-knockout mouse models.

However, the liver possesses, beyond hepatocyte plates, a complex histological and functional architecture that supports metabolic, detoxification, immune, and synthetic activities. Hexagonal-shaped liver lobules are arranged radially with hepatocytes surrounding vascular pericentral or portal structures and lined by liver sinusoidal endothelial cells (LSECs). Within these sinusoids are macrophage-like Kupffer cells (KCs), while hepatic stellate cells (HSCs) are located in the space of Disse. From a pathological perspective, HSCs are a central driver of liver fibrotic disorders, as they constitute the primary source of extracellular matrix (ECM) deposition during fibrogenesis [6, 7]. The anti-fibrotic effects of obeticholic acid (OCA)[8], a semi-synthetic FXR agonist, have led to the idea that FXR is expressed and plays an important role in hepatic stellate cells (HSCs). However, inconsistent reports have failed to agree on FXR expression and anti-fibrotic activities in HSCs [9-13]. More recent studies utilizing single cell approaches have renewed interest in investigating FXR activity in HSCs, which has been detected in both mouse and human HSCs, where FXR maintains a quiescent state [14-19].

We first aimed at deconvoluting FXR expressions and functions across different liver cell types. We then interrogated FXR-controlled responses in unchallenged mice, using a combination of short-term in vivo agonist treatment, hepatocyte *Nr1h4* KO mice and liver cell type isolation coupled to transcriptomic analysis. The characterization of the expression and of the respective functions of hepatocyte and hepatic stellate cell FXRα in naïve liver and an ex vivo model of fibrosis underlined the dichotomic nature of liver FXR.

## MATERIALS and METHODS

*A detailed description of Materials and Methods appears in the Supplemental Information Appendix*.

### Animal experimentation

All procedures complied with EU guidelines for animal experimentation and were approved by the Nord-Pas de Calais Ethical Committee (CEEA75). Unless otherwise indicated, male C57BL/6J mice (8–12 weeks old; Charles River) were maintained under standard conditions with ad libitum access to chow and water. For time-restricted feeding experiments, food access was limited to the dark phase (ZT12–ZT24) for 2 weeks. Unless otherwise stated, mice were euthanized at ZT4–6, and tissues were snap-frozen in liquid nitrogen. For diet-induced MASH/fibrosis studies, mice were fed either a control diet or a high-fat, high-carbohydrate, cholesterol-enriched diet for 24 weeks as described previously [20].

### Liver- and intestine-specific FXR knockout mice

Hepatocyte- and intestine-specific FXR knockout mice were generated by crossing FXR^flox/flox^ mice [20] with Alb-Cre [21, 22] or Villin-Cre [23, 24] transgenic mice, respectively.

### Tropifexor treatment

C57BL/6J mice (12 weeks old, n=6/group) received tropifexor (0.5 mg/kg/day) or vehicle at ZT4 for 3 days. Livers were collected at ZT4–5 for cell isolation or snap-frozen.

### CCl_4_ administration

Mice received CCl_4_ (0.5 mL/kg, i.p.) diluted in olive oil either once or three times weekly for 2 weeks. Livers were subsequently processed for cell isolation and sorting as described previously [18].

### Mouse hepatocyte and hepatic stellate cell isolation

Primary mouse HSCs were isolated from C57BL/6 J mice (male, 15–18-week-old, Charles River Laboratories) according to [18]. Briefly, livers were perfused with collagenase, dissociated, and hepatocytes were recovered by low-speed centrifugation. HSCs were isolated from the non-parenchymal fraction by fluorescence-activated cell sorting based on retinol autofluorescence. Cell viability was assessed using Zombie Green staining.

### The human ABOS cohort

The Hôpital Universitaire de Lille (HUL) cohort (ABOS; ClinicalTrials.gov: NCT 01129297) has been described in detail in [25-27]. Briefly, liver needle biopsies were obtained at the time of surgery from 910 obese patients undergoing bariatric surgery and processed for transcriptome analysis using Affymetrix Human Transcriptome Array (HTA) 2.0. Livers were stratified using stringent qualitative and quantitative criteria into “healthy, normal”, “steatotic,” and “MASH” livers. The fibrosis stage was assessed using the Kleiner scoring system and was used to define a propensity score–matched subcohort based on age, sex, body mass index (BMI), diabetic status and statin use (see [28]). Fifty-three patients with MASH and fibrosis (F>2) were thus matched with 53 MASH-only patients (F=0) [29].

### RNAscope

RNAscope assays were performed on paraffin-embedded human liver sections using the RNAscope Multiplex Fluorescent Reagent Kit v2 (Advanced Cell Diagnostics) according to the manufacturer’s instructions. Human *COL1A1* and *NR1H4* transcripts were detected using RNAscope probes Hs-COL1A1-C2 and Hs-NR1H4-C1, respectively. Following fluorescent signal amplification and detection, sections were immunostained for αSMA (Abcam, ab124964). Images were acquired using an Axioscan Z1 slide scanner (Zeiss).

### Cell lines, plasmids and transfection

#### Cell lines

EMS404 cells (Kerafast, #EMS404) are a mouse hepatic stellate cell line derived from the liver of a *Tlr*4^-/-^ C57BL/6 mouse stably expressing human *TLR4* and immortalized with SV40 large T antigen [30]. AML12 mouse hepatocytes were maintained as previously described [31, 32]. Where indicated, cells were synchronized by treatment with dexamethasone (100 nM) for 2 h prior to experimentation.

#### Transfection

EMS404 cells were maintained in DMEM supplemented with 5% fetal calf serum and penicillin/streptomycin. Transfections were performed using Lipofectamine 2000 according to the manufacturer’s instructions. Twenty-four hours after transfection, cells were treated with GW4064 or tropifexor for 24 h in serum-reduced medium containing 0.2% BSA. For TGFβ stimulation, cells were serum-starved for 9 h and subsequently treated with TGFβ (1 ng/mL) for 24 h in medium supplemented with 0.5% fetal calf serum. The following mouse expression plasmids were used: pcDNA3-FXRα1, pcDNA3-FXRα2, pcDNA3-FXRα3, pcDNA3-FXRα4, and pcDNA3-RXRα [32, 33].

### RNA extraction and RT-qPCR

Total RNA was isolated from cultured cells using the NucleoSpin RNA Kit and from frozen liver samples using TRIzol reagent according to the manufacturers’ instructions. RNA quality and concentration were assessed spectrophotometrically. cDNA was synthesized using the High-Capacity cDNA Reverse Transcription Kit (Applied Biosystems®, Thermo Fisher Scientific), according to the manufacturer’s instructions. Quantitative real-time PCR (qPCR) was carried out using PowerTrack™ SYBR™ Green Master Mix (Applied Biosystems) on a QuantStudio™ 3 Real-Time PCR System (Applied Biosystems®, Thermo Fisher Scientific). Gene expression levels were normalized to *Rps28* or *Rplp0* and calculated using the 2^-ΔΔCt^ method [34].

### Transcriptomic analysis

#### Affymetrix microarrays

GeneChip® Whole Transcript (WT) Expression Arrays (Affymetrix®-Thermo Fisher Scientific, #902281) were used in combination with the GeneChip® WT PLUS Reagent Kit (Affymetrix®, Thermo Fisher Scientific, #902280), following the manufacturer’s protocol.

#### RNA sequencing

Following quality control, RNA libraries were prepared using poly(A) selection and sequenced as paired-end 100-bp reads on the DNBSEQ platform (BGI, Shenzhen, China). Reads were quality-filtered, aligned to the mouse genome (mm10) using HISAT2, and quantified using RSEM [35, 36]. Sample collection and data processing for the RNA-seq analysis of human liver biopsies have been described elsewhere [25]. Human liver RNA-seq data generation and processing have been described previously [23]. FXR isoform expression was quantified using RSEM against the human genome (GRCh38) and Ensembl release 105 transcript annotations.

### Protein extraction, western blotting and WES analysis

Total cellular proteins were extracted in lysis buffer containing protease and phosphatase inhibitors. For subcellular fractionation, cytoplasmic, nucleoplasmic, and chromatin-associated fractions were isolated as previously described [37], and chromatin-bound proteins were released by benzonase digestion. Protein concentrations were determined using BCA assays. Proteins were separated by SDS-PAGE, transferred to nitrocellulose or PVDF membranes, and analyzed by immunoblotting using the indicated antibodies and enhanced chemiluminescence detection. Images were acquired using an iBright FL1500 imaging system. Alternatively, protein expression was quantified using the WES capillary immunoassay system (ProteinSimple) according to the manufacturer’s instructions.

### Preparation and culture of precision-cut liver slices

Precision-cut liver slices (200 µm) were prepared from male C57BL/6J mice harvested at ZT5–7 using a vibratome as previously described [38]. PCLS viability was assessed using Per2::Luc reporter mice [39], and circadian bioluminescence was monitored in the presence of luciferin using a KRONOS-DIO luminometer [38].

### Statistical analysis

Unless otherwise stated, statistical analyses were performed using GraphPad Prism (version 9 or later). Data are presented as mean ± standard error of the mean (SEM). At least three independent experimental replicates were performed. For in vitro data, equality of variances was assessed using the F test. Two-group comparisons were performed using an unpaired two-tailed t-test with Welch’s correction. A one-way ANOVA followed by Tukey’s multiple comparisons test (comparing all groups) was used for multiple comparisons involving one variable. When more than one variable was analyzed, a two-way ANOVA followed by Tukey’s multiple comparisons test was applied.

### Data analysis

Differential gene expression analyses were performed from gene-level count matrices using the Omics Playground v. 3.0 (BigOmics Analytics) [40]. VolcaNoseR was used to generate and label Volcano plots [41].

Cell-cell communication analyses were performed with a modified version of NicheNet [42] using a curated mouse-specific CellTalk ligand-receptor database [43] adapted for bulk tissue transcriptomic data. Single-cell RNA-seq datasets were explored using the ISCEBERG R/Shiny application [44] which was used for dimensionality reduction, clustering, cell-type annotation, and differential expression analyses. Circos plots and other graphical outputs were edited in CorelDRAW for figure preparation

### Datasets

Publicly available datasets used in this study included: the transcriptomic analysis of liver FXR knockout mice [22] (EMBL-EBI: E-MTAB-1722), of mouse livers from the amylin MASH model [45] (NCBI-GEO: GSE129389), of mouse livers under time-restricted feeding [18] (NCBI-GEO: GSE223360), of single-cell and single-nucleus from mouse and human livers [16] (NCBI-GEO: GSE192742) and FXR ChIP-seq analysis in control and TCA-treated mouse livers [33] (NCBI-GEO: GSE87866).

Datasets generated as part of the present study included microarray analysis of total mRNA from EMS404 cells overexpressing FXRα1 or FXRα2 and treated or not with GW4064 (NCBI GEO: GSE331297); RNA-seq analysis of total mRNA from vehicle- or tropifexor-treated precision-cut liver slices (PCLS) prepared from wild-type and hepatocyte-specific FXR knockout mice (NCBI GEO: GSE333328); RNA-seq analysis of liver mRNA from wild-type and hepatocyte-specific FXR knockout mice (NCBI GEO: GSE332832); RNA-seq analysis of hepatocyte mRNAs isolated from vehicle- and tropifexor-treated wild-type mice (NCBI GEO: GSE334504); RNA-seq analysis of hepatic stellate cells isolated from vehicle- or tropifexor-treated mice (NCBI GEO: GSE334831), RNA-seq analysis of liver mRNA from wild-type mice treated with vehicle or tropifexor (GEO NCBI: GSE335123), RNA-seq analysis of liver mRNA from wild-type mice treated with vehicle or OCA (GEO NCBI: GSE335333).

## RESULTS

### *Nr1h4* transcripts and FXR protein expression in liver cell types

Previous studies using whole-body or liver-specific FXR knockout mice have primarily highlighted FXR’s role as a metabolic homeostat (reviewed in [2, 3, 46]). Accordingly, monitoring the activation of the FXR-driven metabolic pathway over time through whole liver gene expression patterns [18] showed that this pathway is most active at the fast-to-fed transition (ZT12) and paralleled the activation of the cholesterol biosynthetic pathway (Supp. Figure 1A). Potential liver FXR target genes were then defined based on 2 criteria: (1) significant downregulation in hepatocyte-specific (Alb-CRE^+/-^, FXR^flox/flox^) FXR knockout mice (|fold change (FC)| > 1.2, p < 0.05, [22] and see below), and (2) presence of an FXR-binding site within ±1 Mb of the transcription start site (TSS) [33, 47, 48]. Using this integrated approach, we defined a set of 698 high-confidence putative FXR target transcripts. Temporal expression analysis revealed that most of these genes peaked between ZT8 and ZT14, which coincided with the onset of the active/feeding phase (Supp. Figure 1B, upper panel). Biological term enrichment analysis (BTEA) revealed a significant overrepresentation of genes associated with lipid metabolism (Supp. Figure 1B, bottom panel). Hepatic FXR protein levels did not vary significantly throughout the day/night cycle (Supp. Figure 1C) and liver primary BA content (cholic acid, CA and the potent natural FXR agonist chenodeoxycholic acid, CDCA; taken from [49]) showed maximal accumulation within the first hours of this active (feeding) period (ZT12-14, Supp. Figure 1D).

While examining the outcome of the selective ablation of *Nr1h4* expression in hepatocytes, we observed that FXR target genes exhibited unequal sensitivity to hepatocyte FXR ablation (Supp. Figure 1E, Supp Table 1). Notably, these genes are significantly expressed in distinct cell types such as *Retsat* [HCs > cholangiocytes (CHs) > KCs > HSCs >L SECs], *Rarres2* (HCs = HSCs) or *Ces1g* (HCs>CHs) ([18]and Supp. Table 1). This graded sensitivity thus suggested that FXR target genes may either function within a broader regulatory network [48], and/or hints at an unaltered FXR-controlled expression in other cell types. To explore the latter hypothesis, we assayed *Nr1h4* transcripts in mouse liver cell types. They were expressed in purified HCs and HSCs in the wild type background, in line with our previous report [18]. In the Alb-CRE^+/-^, FXR^flox/flox^ genetic background, *Nr1h4* transcript expression was maintained in HSCs, whereas it was markedly downregulated in HCs (Supp. Figure 1F).

Controversial studies have accumulated since the seminal paper by Fiorucci and colleagues [9] regarding the expression and roles of FXR in activated hepatic stellate cells (HSCs) [10-12, 50, 51]. We therefore first assayed the expression of FXR in both parenchymal and non-parenchymal liver cells collected at ZT3-5. Using a robust mouse liver cell type sorting procedure[18], significant amounts of highly purified HCs, HSCs, KCs, LSECs and CHs were obtained. *Nr1h4* (encoding FXRα), *Nr1h5* (encoding the mouse-specific FXRβ), *Nr2b1, 2* and *3* (encoding the FXR dimerization partners RXRα, β and γ respectively) transcript levels were quantified by RNA-seq (Figure 1A). *Nr2a1* (encoding HNF4α) was used as a specific HC marker. This analysis revealed that *Nr1h4* expression was restricted to HCs and HSCs, which mostly express *Nr2b1* (a.k.a. *Rxrα*) as the FXR dimerization partner. Examination of transcript structures suggested that HCs expressed the *Nr1h4* α3 and α4 isoforms, whereas HSCs expressed the *Nr1h4* α1 and α2 isoforms. These results were validated by RT-qPCR and confirmed the RNA-seq data (Figure 1B-D). FXR protein (Figure 1E) expression was further quantified by capillary immunoassay and established that HCs and HSCs are indeed the main sites of expression of FXRα in mouse liver (Figure 1F), as expected from single cell (sc)RNA-seq analysis ([15, 16, 52, 53] and Supp. Figure 2A, B). Basal expression levels of genes relevant to BA metabolism demonstrated that HCs are primarily involved in this process, whereas HSCs were essentially devoid of BA-related genes (Figure 1G). Taken together, our data show that murine HCs and HSCs express different FXR isoforms which likely fulfill only partially overlapping functions.

**Figure 1:**
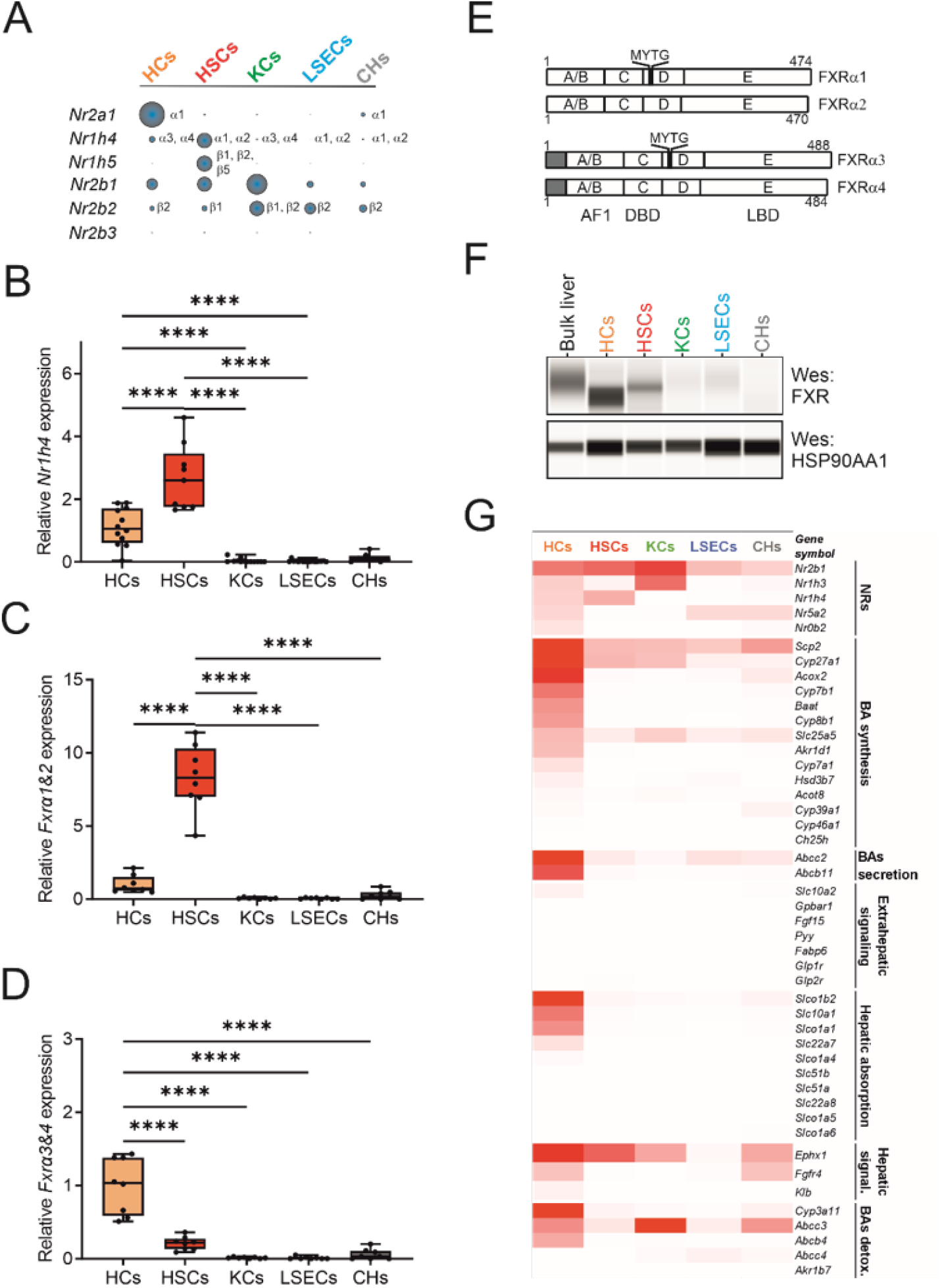
*Nr1h4* gene expression in parenchymal and non-parenchymal cells from mouse liver. A) Whole-transcript and isoform expression of *Nr1h4, Nr1h5, Nr2b1, Nr2b2*, and *Nr2b3*, encoding FXRα, FXRβ, RXRα, RXRβ, and RXRγ, respectively. *Nr2a1*, encoding HNF4α, was used as a hepatocyte marker. Transcripts were quantified by RNA-seq (RPKM), and isoform usage was assessed using Sashimi plots in IGV. B) Quantitative representation of *Nr1h4* expression across liver cell types. Expression levels were determined by RT-qPCR using a primer set amplifying the four mouse *Nr1h4* isoforms. Difference among means (n=8-12) were compared using a two-way ANOVA followed by a Tukey’s multiple comparison test. (p<0.001: ****). C, D) *Nr1h4* isoform expression in liver cell types. Expression levels were determined by RT-qPCR using primer sets specific for the α1/α2 or the α3/α4 isoforms. Difference among means (n=8-12) were compared using a two-way ANOVA followed by a Tukey’s multiple comparison test. (p<0.001: ****). E) Schematic representation of FXR protein isoforms. F) FXR protein expression in liver cell types. G) Basal expression of known FXR target genes. A heatmap of *Nr1h4* and of target genes involved in BA homeostasis and signaling is shown in mouse liver cell types.

### *NR1H4* FXR transcripts and FXR protein expression in human primary hepatocytes and hepatic stellate cells

A similar analysis was performed in human livers. A previous scRNA-seq-based report [16] identified HCs and HSCs as the main site of *NR1H4* expression in human liver (Supp. Figure 3A, B). Quantification of the full-length *NR1H4* transcript by RT-qPCR, performed on purified human HCs and HSCs at very low passages, revealed that *NR1H4* expression occurred predominantly in HCs, and to a lesser extent in HSCs (Supp. Figure 3C). Notably, *NR1H4* transcripts rapidly declined with increasing primary HSC passages (Supp. Figure 3D). *NR1H4* transcripts colocalized in cells expressing *ACTA2* and *COL1A1* (Supp. Figure 3E), and the FXR protein was detected in primary HSCs (Supp. Figure 3F).

Eight *NR1H4* splice variants have been described in primary human hepatocytes [54] and were indeed detected in whole human liver transcripts from non-fibrotic patients ([25], Supp. Figure 4A), including non-functional variants such as the nonsense-mediated decay (NMD) variant and those encoding for truncated versions of FXR (FXRα5-7, [54]). RT-PCR-based detection of human *NR1H4* variants from cultured primary human HCs and HSCs detected the expression of the α1 and α2 isoforms in both HC and HSCs, with minute amounts of α3 and α4 being detected in HCs (Supp. Figure 4B). While revealing some similarities with mouse liver, this analysis was severely hampered by individual variations and the observed rapid decrease of *NR1H4* transcript abundance upon passaging of human primary HSCs, a feature which may partly explain the controversial previously reported results [11].

### Differential FXR isoform transcriptional activity in a mouse HSC cell line

Having established that FXR is expressed in mouse and human HSCs, we characterized the mouse HSC cell line EMS404 [30]. This cell line recapitulates many features of partially activated HSCs [29]. EMS404 cells express low amount of FXR compared to freshly isolated mouse HCs or HSCs or to the murine hepatocyte cell line AML12 [31], while expressing similar levels of RXRα (Figure 2A). EMS404 cells were thus transfected with expression vectors encoding mouse FXRα1, 2, 3 or 4 and mouse RXRα (Figure 2B) to assess the responsiveness of typical FXR target genes to the reference synthetic, non-steroidal FXR agonist GW4064 [55](Figure 2C). The expression of target genes was strongly influenced by the specific FXR isoform which was overexpressed in EMS404 cells. Interestingly, FXRα3 and 4, which we found to be restricted to HCs, were unable to activate the FXR-driven transporters *Abcb11*/*Bsep, Abcb4*/*Mdr2, Nr0b2*/*Shp* or *Rarres2*/*chemerin*. While *Abcb11* and *Abcb4* were equally activated by the α1 and α2 isoforms, *Rarres2* was preferentially activated by FXRα1. In contrast, *Nr0b2* was only sensitive to the ligand-activated FXRα2 isoform (Figure 2C). This isoform selectivity was further explored by a whole transcriptome analysis of FXRα1 or FXRα2-transfected EMS404 cells (Supp. Figure 5). This approach confirmed that subsets of agonist-activated genes were equally sensitive to α1 and α2 isoforms (such as *Osgin1* and *Fabp6*) or preferentially activated by the α1 isoform (such as *Gsta2* and *Gstm1*) or solely by the α2 isoform (such as *Aqp1* or *Slc03a1/Oatp*). Therefore, FXRα1 and α2 isoforms are preferentially expressed in HSCs in which they may exert distinct regulatory roles.

**Figure 2:**
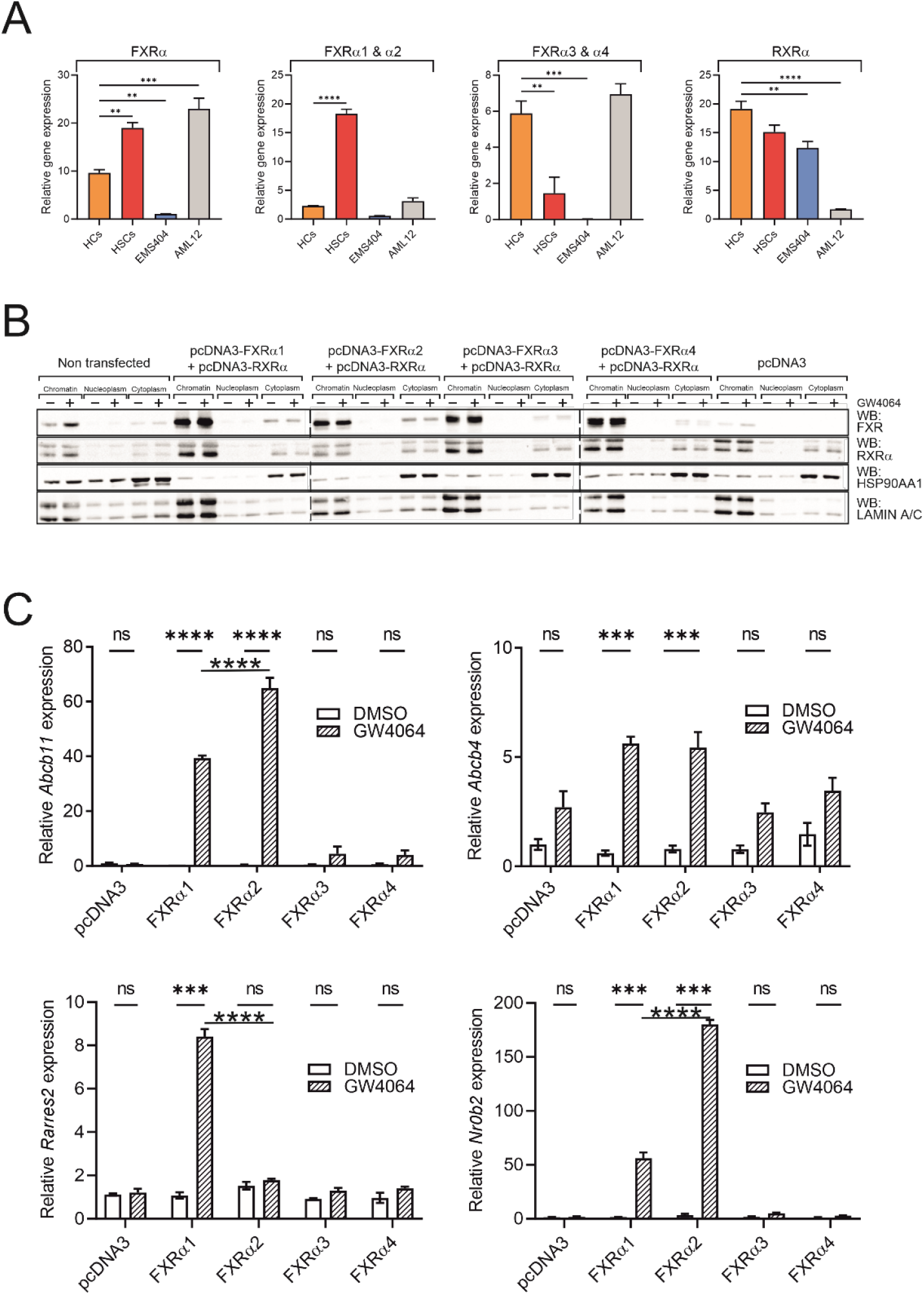
Gene selectivity of FXR isoforms in the HSC cell line EMS404. A) Relative expression level of *Nr1h4* in primary cells and cultured cell lines. RNA from mouse primary hepatocytes (HCs), mouse primary hepatic stellate cells (HSCs) and from EMS404 and AML12 cell lines were analyzed by RT-qPCR to quantify the indicated *Nr1h4* transcript isoforms. Expression of the transcript encoding its obligate heterodimerization partner, RXRα, was also measured. Difference among means (n=3) were compared using a one-way ANOVA followed by a Tukey’s multiple comparison test (p<0.01: **; p<0.005: ***; p<0.001: ****). B) Quantification and subcellular localization of FXRα and RXRα in transfected EMS404 cells. After transfection, cell extracts were collected and fractionated into cytosolic, nuclear, and chromatin-bound fractions. FXRα, RXRα, and subcellular compartment markers (HSP90AA1 and lamin A/C) were detected by WES analysis. C) Selective gene activation by FXR isoforms. EMS404 cells were transfected with the indicated expression vector and treated or not by GW4064 overnight. Transcripts were quantified by RT-qPCR. Difference among means (n=3) were compared using a one-way ANOVA followed by a Tukey’s multiple comparison test (p<0.005: ***; p<0.001: ****).

### Comparative mouse liver transcriptomic response to bile acids and non-bile acid FXR agonists

The FXR signaling pathway can be pharmacologically activated using (semi)synthetic agonists, whose activity has been assessed in pathological models [45, 56, 57] with potential applications in human MAFLD and fibrosis [58]. In addition to these therapeutic actions, FXR agonist treatment can also mimic, in healthy mice, the acute buildup of bile acids such as it occurs in cholestasis or liver regeneration [59, 60].

To identify the most effective agonist active in healthy mice, we analyzed the liver’s transcriptional response after a 3-day administration of obeticholic acid (OCA) (i.p., 30 mpk/day) in comparison to the isoxazole derivative tropifexor (i.p., 0.5 mpk/day), an improved version of GW4064 (Supp. Figure 6). Consistent with a previous report [45], tropifexor triggered a broader transcriptional response compared to OCA (Supp. Figure 6A), upregulating 2,242 genes under these conditions, while OCA upregulated only 114 genes, 75% of which were also regulated by tropifexor (Supp. Figure 6B, Supp. Table 2). Evidence of target engagement by tropifexor was provided by the strong induction of classic FXR target genes such as *Akr1b7* (log2FC=7), *Fabp6* (log2FC=5.4), and *Fmo3* (log2FC=7) (Supp. Table 2), along with the downregulation of the bile acid (BA) biosynthetic pathway (Supp. Figure 7A). BTEA analysis against the Gene Ontology Biological Process (GO BP) database showed that both FXR agonists predominantly induced a shared mitogenic response (Supp. Figure 6B, C).

Exogenous BAs can prime a mitogenic program in the non-injured liver, a hepatocyte FXR-dependent response that lowers the threshold for hepatocyte cell-cycle entry in regenerative contexts [61, 62]. Liver growth in such regenerative settings is also governed by intestinal FXR signaling through FGF15 [63, 64]. We therefore assessed whether intestinal FXR signaling contributes to the observed hepatic mitogenic response under unchallenged conditions. Upon administration of tropifexor to intestinal FXR-deficient mice, we observed that hepatic, cell cycle-related genes such as *Foxm1* and *Mki67* were still significantly upregulated in this genetic background (Supp. Fig. 7B). Thus, the hepatic FXR-driven mitogenic response in normal livers occurs independently of intestinal FXR signaling.

Interestingly, BTEA of tropifexor-upregulated genes, excluding cell cycle-related genes, revealed enrichment in pathways related to cell migration, shape, and polarity. A closer look at the list of genes upregulated by tropifexor (Supp. Table 2) also confirmed the reported induction of an antioxidative response via the coordinated induction of GST-encoding genes [45]. Overall, these findings suggest that supra-physiological levels of BA analogs in healthy mice trigger a liver mitogenic response, suppress BA synthesis, and promote anti-oxidative responses, thus mimicking certain early-stage responses to liver injury.

### Comparative hepatocyte and hepatic stellate cell transcriptomic response to FXR agonism

To differentiate the hepatic response to tropifexor stemming from HCs or HSCs under these conditions, we isolated each liver cell type after a 3-day treatment and then conducted RNA-seq analysis. The 2,392 upregulated genes in HCs as well as the 983 upregulated genes in HSCs were strongly enriched in genes involved in cell cycle/division (Figure 3A, B). Repressed genes (n=1,670) in HCs belonged mostly to lipid metabolic pathways, whereas BTEA barely identified significantly enriched pathways in HSCs from the 389 downregulated genes. Comparing associated functions of commonly tropifexor-modulated transcripts identified a set of upregulated genes (n=378) enriched in cell cycle/division-related transcripts (Figure 3C). Upregulated HC-specific transcripts (n=2,014) were enriched in transcripts involved in cell migration and cytoskeleton organization, whereas HSC-specific transcripts were preferentially annotated with terms related to immune responses. Downregulated transcripts showed a strong cell type specificity, with only 2.4% of shared transcripts with no associated functions (Figure 3C).

**Figure 3:**
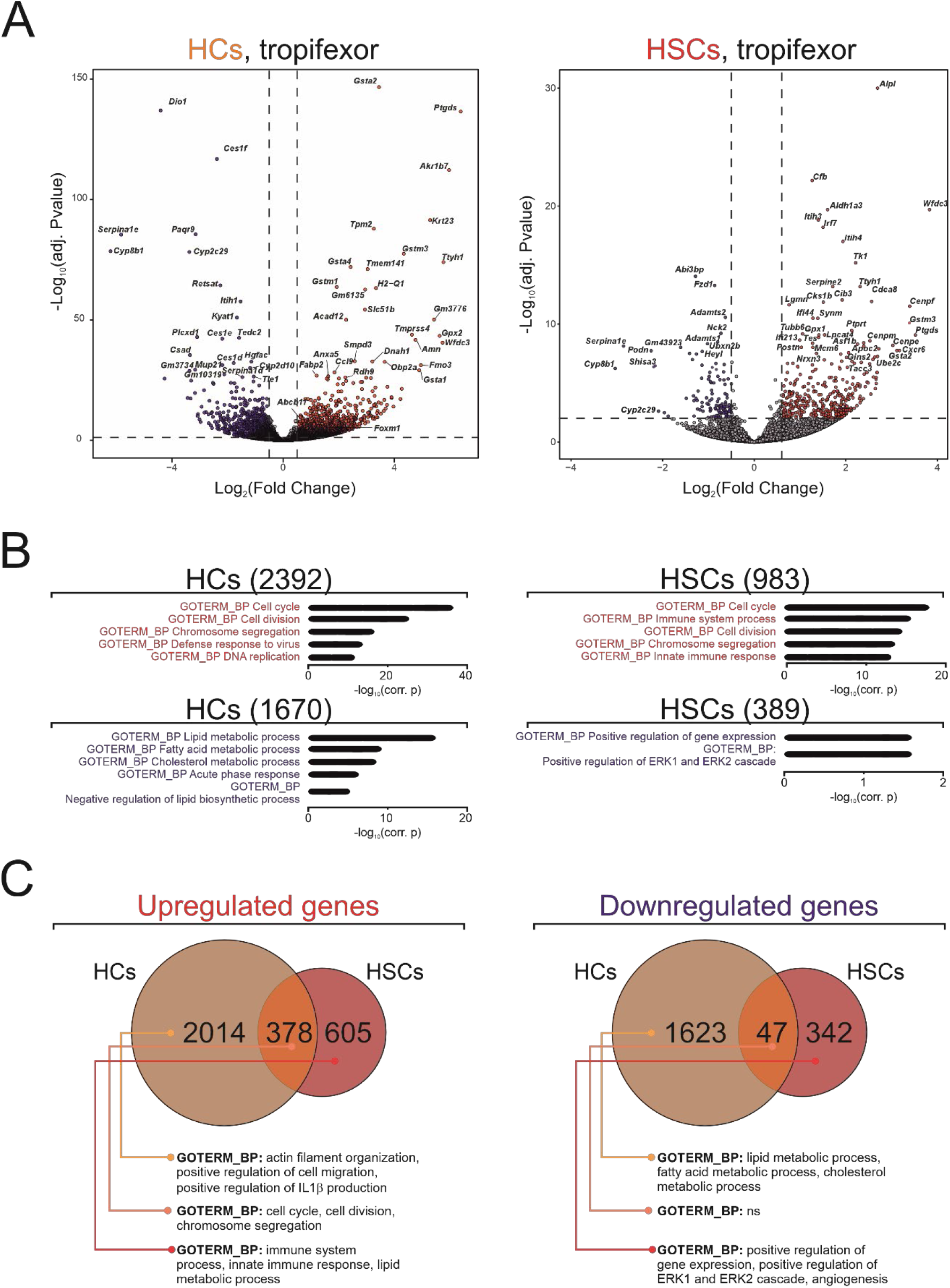
Comparative transcriptome of hepatocytes and hepatic stellate cells from tropifexor-treated mice. A) Volcano plots of differentially expressed genes in HCs and HSCs isolated from livers of mice treated with tropifexor (0.5 mpk) or vehicle for 3 days. Red: upregulated, blue downregulated. B) Biological term enrichment analysis of up- or downregulated genes upon in vivo tropifexor treatment in HCs and HSCs. Red: upregulated, blue downregulated. C) Venn diagrams of differentially up- or downregulated genes in HCs and HSCs and corresponding BTEA analysis. The 3 top hits in the Biological Process database are shown.

The cell cycle-related response was broader in HCs than in HSCs (185 associated transcripts in HCs vs 96 in HSCs)(Supp. Figure 8). However, 85% of HSC-expressed genes were also upregulated in HCs, suggesting a common tropifexor-induced proliferative response in both cell types. Tropifexor-exposed HCs expressed a specific 103-gene subset which, based on further BTEA and GSEA analysis, was not annotated with terms other than those relating to cell cycle and cell division. Using the ChEA EnrichR database which integrates data from ChIP-based experiments [65], FOXM1 was identified as a likely transcriptional regulator of common and HC-specific cell cycle-related upregulated genes (Supp. Figure 8). This aligns with the general properties of FOXM1 as an FXR-regulated cell cycle activator in liver [61, 66]. However, a similar analysis of HSC-specific, cell cycle-related genes rather suggested that MYC, whose transient overexpression precedes HSC activation [67], may exert specific regulatory effects in HSCs on these genes.

Potential direct FXR target genes were identified based on their induction by FXR agonists and the occurrence of 1 or several FXR binding sites in the vicinity of their transcription start site (TSS). More than 80% of tropifexor-induced genes displayed at least 1 cis-regulatory FXR chromatin binding site within 1 Mb of their TSS (Supp. Figure 9A). A GO BP-based BTEA as performed above yielded a similar enrichment for cell type-specific and commonly upregulated genes, showing that observed mitogenic and other responses are very likely resulting from FXR activation. To further illustrate FXR-specific functions in HCs or HSCs, we identified transcripts specifically expressed in each cell type and upregulated by tropifexor. Only a few such genes were identified: 16 in HCs and 12 in HSCs (Supp. Table 3). Among these, *Serpina5* (a serine protease inhibitor) and *Scube1* (a secreted BMP antagonist with putative liver protective activity [68]) were selected as HC- and HSC-specific transcripts, respectively (Supp. Figure 9B). Both loci showed multiple FXR binding sites and *Serpina5* and *Scube1* transcripts increased significantly upon tropifexor administration in a cell type-dependent manner.

This analysis was extended to GST-encoding genes, whose anti-oxidative activities could be related to the protective effect of tropifexor in MASH models [45]. In non-stimulated liver cells, GST-encoding transcripts were mostly detected in HCs and albeit at lower levels, in HSCs and CHs (Supp. Figure 10A). RNA-seq analysis of bulk liver tissue revealed significant upregulation of these genes following tropifexor treatment. (Supp. Figure 10B). Comparison of the transcriptional response into HC- and HSC-specific components showed that *GSTa4, a5, m2, m4, m2, p2, t1, t2, t3* were selectively induced in HCs, whereas *GSTa3* and *m6* were solely induced in HSCs. Interestingly, GSTα3 has been shown to prevent HSC activation and fibrogenesis [69]. Others GST isotypes (*a1, a2, m1, m3, p3*) were equally induced in both cell types (Supp. Figure 10B). All GST-encoding loci harbored FXR binding sites in the vicinity of their TSS, as shown by ChIP-Seq analysis of bulk hepatic tissue (Supp. Figure 10C).

Intercellular communication (ICC) is also key to liver homeostasis but the role of FXR as an ICC regulator has not been explored. The regulation of HC and HSC ligand (L)-receptor (R) pairs harboring FXR binding sites was therefore examined using the CellTalk mouse database [43](Supp. Table 4). BMP, collagen and semaphorin family members were amongst functionally important components of this cellular interactome to be upregulated by tropifexor specifically in HCs (162 total), while integrins and interleukin receptors were detected as such in HSCs (47 total). Genes encoding for chemokines (*Ccl2, 4, 5, 9*), *Spp1*/*osteopontin* and *Rarres2*/chemerin were upregulated in both cell types.

Taken together, this data suggests that FXR has specific roles in HC and HSC that affect both cell-autonomous functions and liver ICC.

### FXR agonism alleviates the fibrotic response in an ex vivo model

Systemic FXR agonism has beneficial effects on MASH and fibrosis progression in multiple rodent models [45, 56, 70] and in humans [58]. The mechanistic basis for these effects is still ill-defined but have been mostly ascribed to metabolic effects and interference with the inflammasome or pro-inflammatory pathways [71]. To further investigate the role of FXR signaling in regulating fibrosis progression, we used the ex vivo precision-cut mouse liver slice (PCLS) model of fibrosis (Figure 4). Under our conditions, PCLS initiated a fibrotic response by day 3 post-isolation (D3), which progressively intensifies through day 9 (D6, Figure 4A). The fibrotic progression was exemplified by the upregulation of genes involved in extracellular matrix (ECM) remodeling, including collagens, TGFβ-related factors and metalloproteinases.

**Figure 4:**
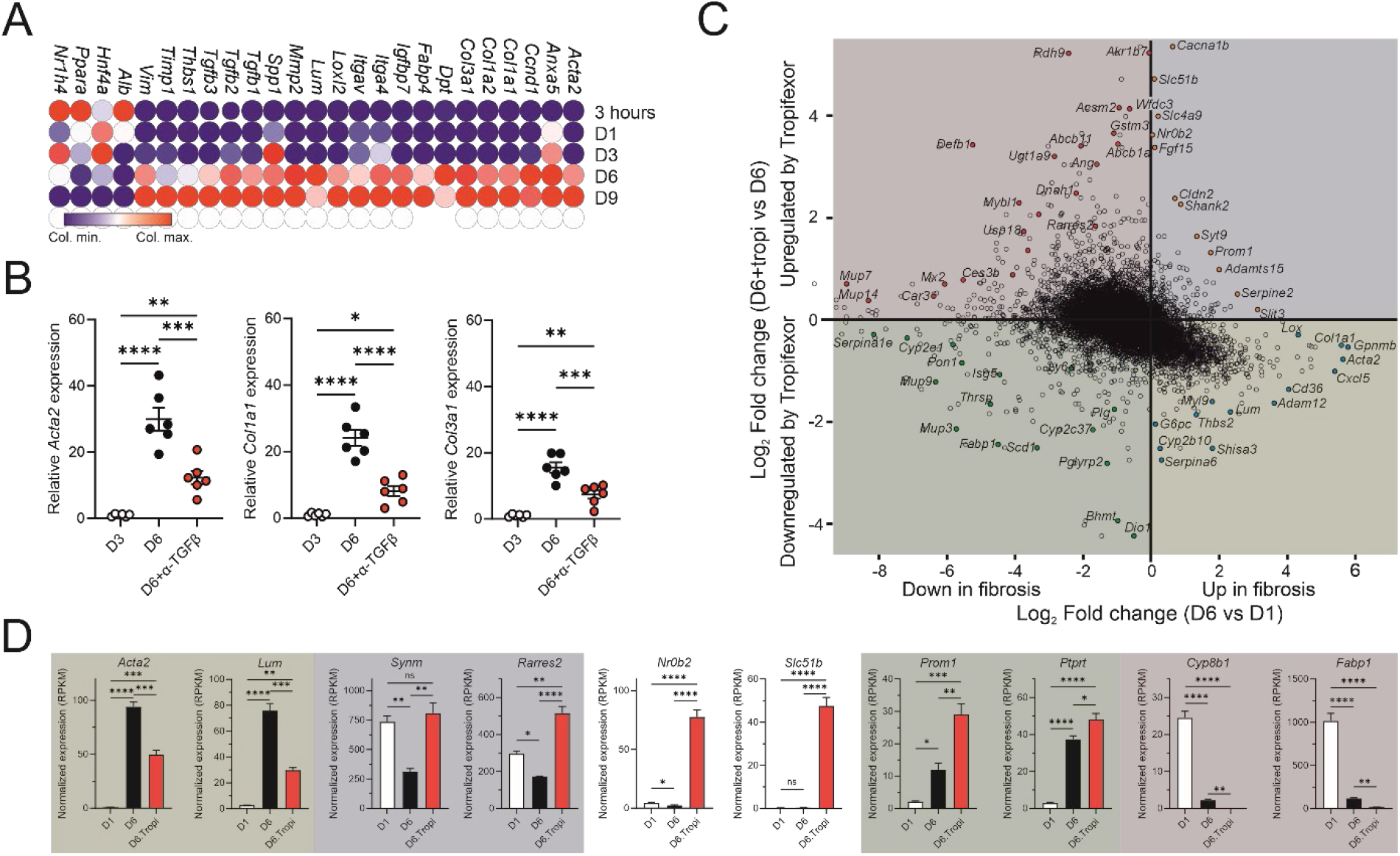
Tropifexor-induced gene expression in mouse precision-cut liver slices. A) PCLS spontaneously acquire a pro-fibrotic transcriptional profile over time. Gene expression was monitored in PCLS from male mice by RNA-seq at the indicated time points [3 hours, one (D1), three (D3) and six days (D6) after preparation]. In the bubble plot, transcript levels are represented as a function of time by a color gradient ranging from blue (lowest expression) to red (highest expression). B) The spontaneous pro-fibrotic response of PCLS is TGFβ-dependent. PCLS were treated with an anti-TGFβ antibody on day 3 (D3) after preparation. Relative expression levels of the indicated genes were measured at the indicated time points. C) Two-dimensional differential expression scatter plot of fibrosis- and tropifexor-regulated genes at D6. PCLS were treated with tropifexor, or left untreated, from D3 to D6, and RNA was extracted at D6. D1 RNA samples were used as the reference to assess fibrosis progression up to D6. Each dot represents a gene positioned according to its log_2_ fold change during fibrosis progression (x axis) and its log_2_ fold change in response to tropifexor treatment at D6 (y axis). D) Selected genes for each quadrant from the scatter plot shown in C). Difference among means (n=3) were compared using a one-way ANOVA followed by a Tukey’s multiple comparison test (p<0.05: *; p<0.01: **; p<0.005: ***; p<0.001: ****).

The fibrotic response was highly dependent on TGFβ signaling, as PCLS treatment with an anti-TGFβ1, 2, 3 antibody markedly suppressed fibrotic gene expression at D6 (Figure 4B). A transcriptome analysis was performed on PCLS undergoing fibrosis (D6) and treated with tropifexor or vehicle (from D3 to D6) (Figure 4C, D). Expression of canonical FXR target genes (*Akr1b7, Slc51b, Nr0b2, Fgf15*) was not affected by fibrosis progression but remained highly responsive to FXR agonism.

Tropifexor efficiently blunted the induction of profibrotic genes such as *Acta2, Lum, Cxcl5, Adam12*. Interestingly, FXR agonism restored the expression of *Synm* and *Nrxn3*, markers of quiescent HSCs, as well as *Rarres2* associated with the inactivated HSC phenotype [72-74]. Additional distinctive expression profiles were identified: *Prom1*/*Cd133*, a fibrosis-protective factor [75, 76], was induced in fibrosis and further increased upon FXR agonism. *Prprt*, encoding a receptor-type protein tyrosine phosphatase acting on STAT3 and associated to HSC quiescence [73], showed a similar expression profile.

*Cyp8b1*, which encodes a cytochrome P450 enzyme responsible for cholic acid synthesis via 12α-hydroxylation, was repressed in fibrosis and further repressed by tropifexor, consistent with previous reports [77]. Similarly, *Fabp1*, involved in intracellular fatty acid transport and whose deletion confers protection from diet-induced hepatic steatosis and fibrosis [78], was similarly regulated.

Taken together, these data suggest that FXR signaling plays a protective role during liver fibrogenesis by blunting profibrotic gene expression, amplifying protective responses and promoting a quiescent or inactivated hepatic stellate cell phenotype. It further supports the potential role of FXR in regulating liver ICC.

### FXR plays a key role in attenuating HSC activation in response to profibrotic stimulus

Since FXR is expressed in both HCs and HSCs, we assessed which of this cell type predominantly contributed to the observed FXR-mediated, anti-fibrotic activity. We generated PCLS from wild type (Alb-CRE^-/-^, FXR^flox/flox^) and hepatocyte *Nr1h4* knockout (FXR^ΔHC^) knockout (Alb-CRE^-/+^, FXR^flox/flox^) mice and first compared liver fibrosis progression in these 2 genetic backgrounds (Figure 5A). No major differences in basal gene expression were observed between the 2 backgrounds, indicating that hepatocyte-specific deletion of FXR does not affect the liver’s sensitivity to fibrosis induced by ex vivo mechanical injury. Differential expression analysis of tropifexor-induced fold changes in both genetic backgrounds revealed distinct patterns of gene responses as illustrated in Figure 5B. Depletion of FXR in hepatocytes impaired the induction of certain genes, including canonical FXR targets (cluster 1). In contrast, cluster 2 genes were similarly upregulated regardless of hepatocyte FXR status. The transrepressive activity of FXR remained intact for cluster 3 genes in both backgrounds, whereas full repression of cluster 4 genes required hepatocyte-derived FXR.

**Figure 5:**
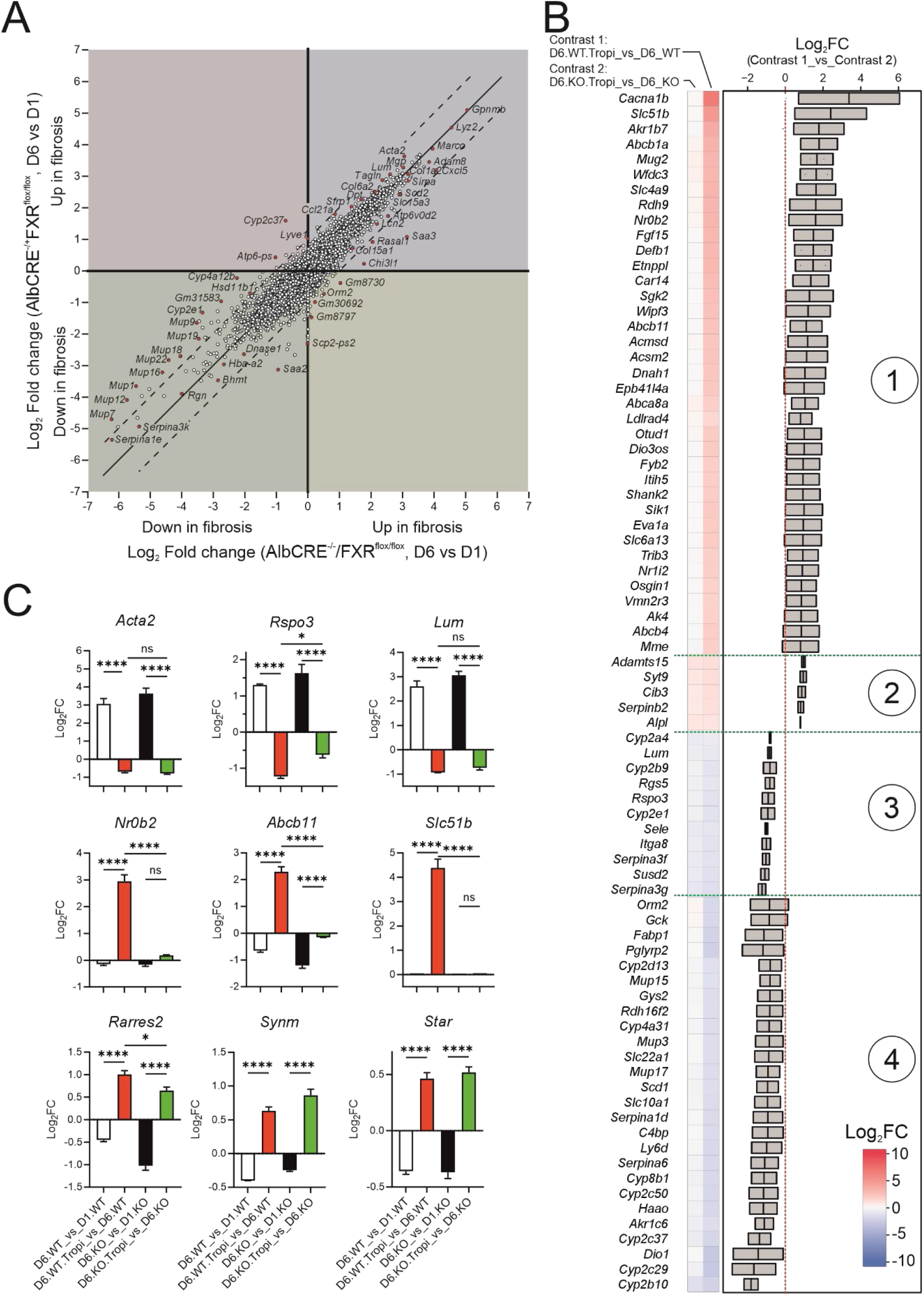
Fibrosis and FXR agonist responsiveness in FXR^ΔHC^ PCLS. A) Two-dimensional differential expression scatter plot of fibrosis-regulated genes at D6 in wild type vs FXR^ΔHC^ mouse PCLS. PCLS from each genetic background were left untreated from D1 to D6, and RNA was extracted at D6. D1 RNA samples were used as the reference to assess fibrosis progression up to D6. Each dot represents a gene positioned according to its log_2_ fold change during fibrosis progression in progression in wild type PCLS (x axis) and its log_2_ fold change during fibrosis progression in FXR^ΔHC^ PCLS (y axis). B) Signature clustering of tropifexor-modulated gene expression in wild type vs FXR^ΔHC^ PCLS. Genes with the most similar expression patterns were grouped into 4 clusters and are displayed as a heatmap (left panel). The relative fold change of selected genes is shown in the right panel. C) Comparative gene expression across all four conditions. Expression of selected genes was validated by RT-qPCR and is shown as log_2_FC, as indicated. Difference among means (n=6) were compared using a two-way ANOVA followed by a Tukey’s multiple comparison test (p<0.05: *; p<0.01: **; p<0.005: ***; p<0.001: ****).

To illustrate the contribution of hepatocyte and HSC FXR (FXR^HC^, FXR^HSC^) to the transrepression of profibrotic genes, we studied the expression values of the HSC-restricted genes *Acta2* (encoding αSMA), *Rpso3* (encoding R-SPONDIN 3) and *Lum* (encoding LUMICAN). Tropifexor treatment similarly blunted the fibrosis-induced overexpression of these 3 representative genes (Figure 5C). A similar analysis was carried out on the canonical, hepatocyte-restricted *Nr0b2, Abcb11* and *Slc51b* genes. The tropifexor-induced expression of these genes was strictly dependent on FXR^HC^ expression. Further analysis of additional relevant transcripts (*Rarres2, Synm, Star*) identified a third category of genes. These genes were repressed in fibrosis, but their expression was similarly restored in both genetic backgrounds by tropifexor, highlighting the key role of FXR^HSC^ in regulating their activity.

Taken together, this analysis suggests that FXR activation normalized aberrant gene expression, including genes with key roles in HSC biology such as *Rarres2* (associated to the inactivated HSC phenotype [73]), synemin (*Synm*) encoding a type of intermediate filament and a marker of quiescent HSCs [72] and R-spondin 3 (*Rspo3*), encoding a potent enhancer of Wnt/β-catenin signaling, whose deletion in HSC significantly protects the liver from pericentral toxicants such as APAP and CCl_4_ [79].

### Chemerin expression is altered in mouse and human fibrosis

To assess how ICC homeostasis is altered in mouse and human fibrosis, we first characterized 2 mouse models for advanced fibrosis to identify active L(igand)-R(eceptor) pairs using a version of the NicheNet predictive tool adapted to the analysis of bulk RNA-seq datasets [42]. We first used a chronic 7-week CCl_4_ treatment model which induces pericentral damage and fibrosis. Up- or downregulated, active L-R pairs were inferred by computational analysis (Supp. Figure 11). Crucial mediator of liver fibrosis such as the chemokine *CCl3*, the tissue inhibitor of metalloproteinases-1 (*Timp1*) or the transforming growth factor β1 (*Tgfb1*) were found, amongst others, to be highly active in profibrotic conditions. In naïve livers, pathways driven by the protective chemokines *Cxcl12*/*Sdf1* and *Rarres2*/chemerin, and the key determinant of resistance to liver injury *Col18a1* were active. A similar analysis was applied to a diet-induced MASH and fibrosis model [20]. Activated and repressed pathways were distinct from those engaged in the chemically induced injury model, but quite strikingly also identified the *Rarres2* pathway as significantly downregulated in this diet-induced MASH model (Supp. Figure 12). Taken together, these analyses thus showed that the downregulation of the *Rarres2* pathway is a common feature of advanced MASH and of fibrosis.

Plausible causes for this fibrosis-induced downregulation were investigated. FXR expression was primarily reduced in hepatocytes (HCs), but not in hepatic stellate cells (HSCs) (Supp. Figure 13A, B), and *Nr1h4* mRNA splicing remained unaffected in CCl_4_-challenged livers (Supp. Figure 13C, D). Transcripts encoding preprochemerin were reduced in human steatotic and fibrotic livers (Supp. Figure 14). *NR1H4* transcript levels were barely affected in MASH patients but not in fibrotic patients, and splicing was unchanged in those 2 conditions (Supp. Figure 15). *Rarres2* transcripts were also significantly upregulated by tropifexor in a second insulin-resistant, obese mouse model of MASH and fibrosis (Amylin [AMLN] diet, [45])(Supp. Figure 14C), in line with our in vitro data.

### FXR agonism recapitulates the antifibrotic benefits of chemerin pathway blockade

We thus designated *Rarres2* as a direct FXR target gene based on 2 observations showing that (i) FXR binds 3’ of the *Rarres2* locus in mouse liver (Supp. Figure 16) and (ii) *Rarres2* expression is mostly driven by the HSC-restricted FXRα1 isoform (Figure 1A, 3C) whose expression is not significantly altered in fibrotic D6 PCLS (Supp. Figure 17). Given that FXR^HC^ ablation only partially diminished tropifexor-induced *Rarres2* expression (Figure 5), this further suggested that HSC-derived chemerin may play a key role in the protective effects of FXR against fibrosis.

To gain mechanistic insight into chemerin signaling in healthy and fibrotic mouse livers and mouse PCLS, we systematically profiled the expression of key components of its pathway. They include *Rarres2*, proteases mediating its C-terminal cleavage and its extracellular activation or inactivation [80, 81], and its cognate receptors (*Cmklr1, Gpr1, Ccrl2* [82])(Figure 6). In unchallenged mouse liver, protease-encoding transcripts were predominantly expressed in hepatocytes (HCs), including *Plg* (encoding plasminogen). Activation of plasminogen requires cleavage by urokinase-type plasminogen activator (*Plau*), expressed in Kupffer cells (KCs), or by coagulation factor XII (*F12*), which is expressed in HCs (Figure 6A). Carboxypeptidases B2 and N2 (*Cpb2, Cpn2*), which also contribute to chemerin maturation, were mainly detected in HCs. Cathepsins L and S (*Ctsl, Ctss*) were expressed across multiple cell types but were most abundant in HCs, KCs, and CHs. Finally, mRNAs encoding the signal-transducing chemerin receptor CMKLR1 were almost exclusively detected in KCs. Taken together, these gene expression profiles suggest that chemerin signaling in the mouse liver may function through a paracrine axis, originating from HCs and HSCs and targeting KCs.

**Figure 6.**
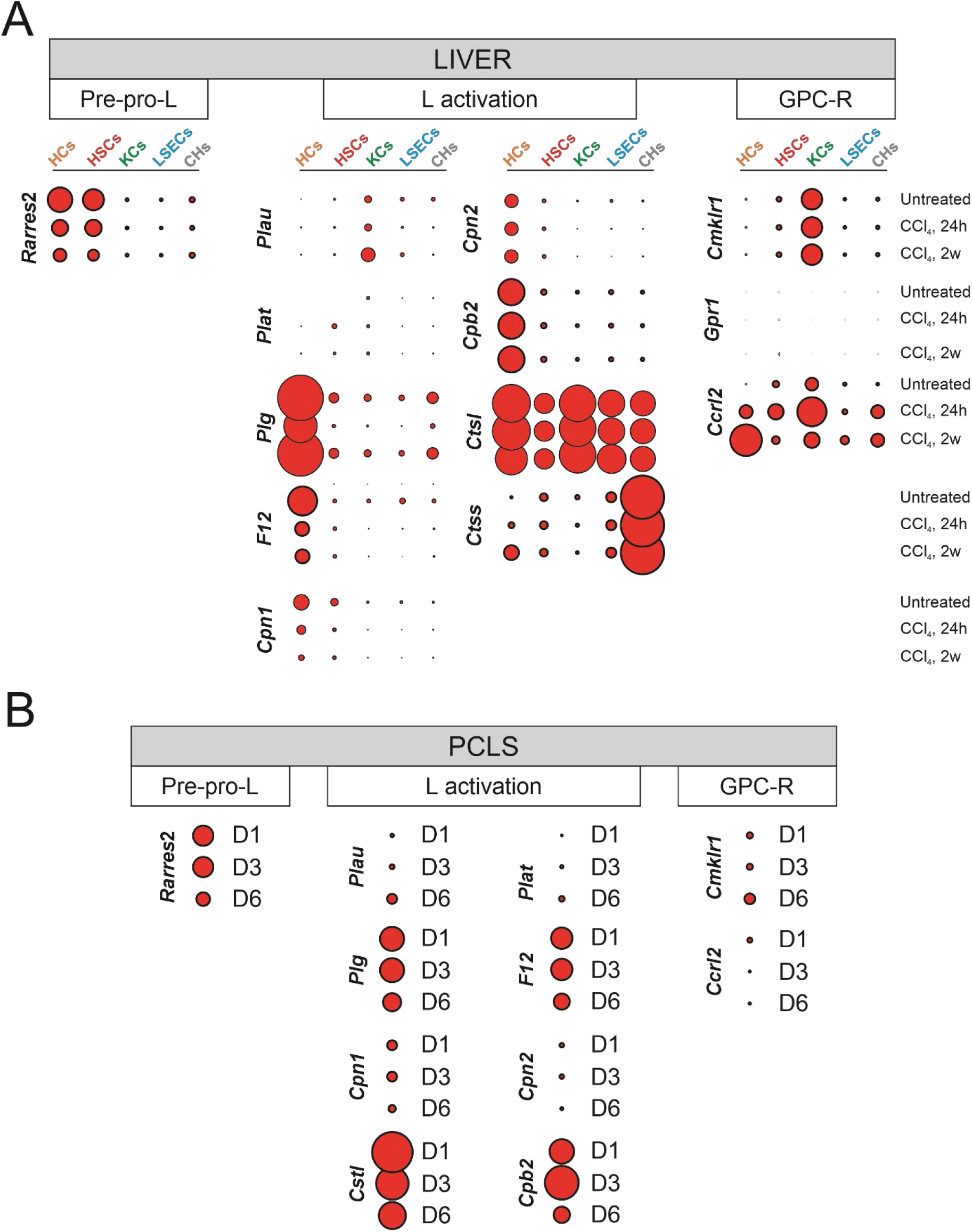
The chemerin pathway in mouse liver cell types and PCLS. A) Expression of genes encoding components of the chemerin pathway in liver cell types. HCs, HSCs, KCs, LSECs, and CHs were isolated from male mouse livers exposed to vehicle or CCl_4_ for 24 hours or 2 weeks, and their transcriptomes were analyzed by RNA-seq. Expression values for the indicated genes are shown as a bubble plot, with bubble size proportional to RPKM values (Red: upregulated). B) Expression of genes encoding components of the chemerin pathway in mouse PCLS. Transcriptomes from PCLS were analyzed by RNA-seq at the indicated time points (1, 3 or 6 days after preparation, denoted D1, D3, D6). Expression values for genes encoding components of the chemerin pathway are shown as above.

Interestingly, a human liver spatial reference atlas has recently been generated [83]. Analysis of the concordant zonation for *RARRES2* and its cognate receptors suggested that zone-specific intercellular communication most likely occurs between fibroblasts, hepatocytes and Kupffer cells (Supp. Figure 18), consistent with our mouse data.

Treatment by the fibrosis inducer CCl_4_ did not alter the expression of most of these components, with the exception, as reported above, of *Rarres2* itself, *F12* and *Cpn1* (Figure 6A). In PCLS, gene expression profiles followed a similar general trend (Figure 6B), thus validating this ex vivo model for further functional studies. FXR agonism restored *Rarres2* expression in a profibrotic milieu (Figures 4D, 5C, Supp. Figures 11, 12). We therefore examined whether the chemerin pathway is at play during the fibrosis response observed in PCLS. Since commercially available chemerin batches contained multiple cleaved peptides hence low amounts of full length chemerin, we treated PCLS (from D3 to D6) with the biased peptidic agonist C9 [84], which preferentially activates the GPCR branch downstream of CMKLR1 but not the β-arrestin pathway [85]. C9 treatment did not reduce the expression of fibrosis markers *Acta2, Col1a1*, and *Col3a1* (Figure 7A), suggesting either a degradation of this short peptide in PCLS or its inability to displace CMKLR1-bound chemerin [86]. We thus treated PCLS with the chemically more stable CMKLR1 inhibitor α-naphthylethyltrimethylammonium iodide (α-NETA), which efficiently blocks both GPCR- and β-arrestin-coupled pathways [87]. In contrast to C9, α-NETA suppressed the induction of fibrosis markers as effectively as tropifexor, but neither showed synergistic effects when used together (Figure 7B). This suggests that tropifexor-modulated chemerin might inhibit the CMKLR1 pathway either by being processed into antagonistic forms through abnormal cleavage or by promoting CMKLR1 desensitization [81].

**Figure 7:**
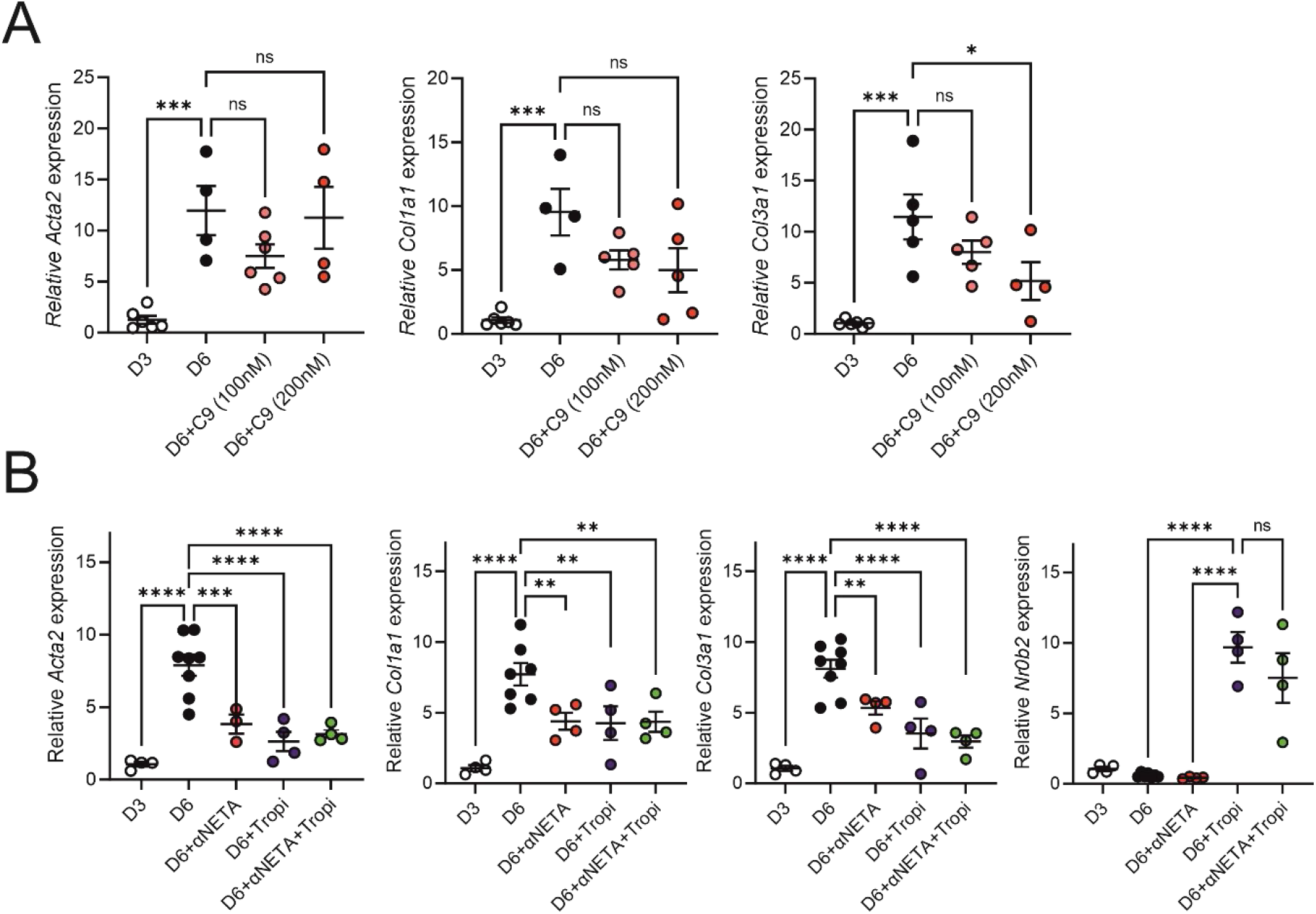
Modulation of the chemerin pathway during fibrosis progression ex vivo. A) Chemerin-9 (C9) activity in PCLS. PCLS were prepared as above and treated 3 days later (D3) with increasing doses of the C9 nonapeptide. RNAs were extracted 3 days later (D6) to characterize by RT-qPCR the expression of representative profibrotic genes. Difference among means (n=4-6) were compared using a one-way ANOVA followed by a Tukey’s multiple comparison test (* p<0.05;*** p<0.005). B) αNETA activity in PCLS. PCLS were prepared as above and treated 3 days later (D3) with αNETA and/or tropifexor. RNAs were extracted 3 days later (D6) to characterize by RT-qPCR the expression of representative profibrotic genes. Difference among means (n=4-6) were compared using a one-way ANOVA followed by a Tukey’s multiple comparison test (** p<0.01; **** p<0.001).

## DISCUSSION

The nuclear bile acid receptor FXRα gained prominence through the characterization of its role as a bile acid sensor and of its metabolic regulatory functions along the intestine-liver axis. Early studies also highlighted its involvement in liver regeneration [61, 66] and, more generally, in hepatoprotection [59]. FXRα thus emerges as a pleiotropic regulator operating across several organs, cell types, and pathophysiological conditions. Importantly, this functional diversity is also influenced by its primary structure, as FXRα is expressed as several isoforms with distinct biological properties, as shown in hepatocytes [88]. Deciphering the processes that govern FXRα activity in resident liver cell types is therefore a crucial first step toward fully understanding FXR pathophysiology and harnessing the therapeutic potential of (semi-)synthetic FXR agonists to correct liver dysfunctions.

Here, using a carefully controlled liver cell-sorting procedure [18], we show that HSCs express substantial amounts of FXR, a finding further supported by single cell/nucleus RNA sequencing data. We also show that FXRα isoforms are differentially expressed in HCs and HSCs, with HCs predominantly expressing the α3/α4 isoforms and HSCs mainly expressing the α1/α2 isoforms. In vitro functional characterization of FXRα isoforms in a relevant mouse HSC cell line revealed distinct capacities of the different isoforms to activate FXR target genes, raising the possibility that an isoform switch within the HSC population may influence cellular responsiveness to bile acids. Isoform-selective transcriptional activity has previously been described in a hepatocyte background [88-91]. However, little is known about the mechanisms regulating isoform switching, although fasting and exercise have been proposed as physiological cues associated with increased FXRα2 expression [92]. Because fibrosis progression did not measurably alter isoform balance in our mouse models, the conditions and mechanisms required for a biologically meaningful isoform switch in these cells remain to be established.

Gene expression analysis in unchallenged livers showed that processes involved in maintaining bile acid homeostasis are largely confined to HCs. To further investigate the contribution of FXR to HSC biology, we analyzed the transcriptome of HSCs and of HCs exposed in vivo to OCA or to the FXR-selective agonist tropifexor. We identified a marked cell cycle-related response, which was also detected in HCs. This response was independent of intestinal FXR expression and resembles that observed in naïve liver following a cholic acid-enriched diet [61]. HC proliferation is an essential component of liver regeneration and hepatoprotection, and whole body knock out of *Nr1h4* severely impairs liver regeneration [61]. The role of FXR^HC^ in these processes is likely to be ancillary to growth factors like EGF and PDGF [59], but our data suggest that FXR activation in HSCs might also participate in this response. Consistent with this idea, HSCs have recently been shown to support homeostatic hepatocyte proliferation by forming mitogenic niches [93] which might expand in regenerating liver and further contribute to re-establishment of liver zonation [79]. Intriguingly, we did not detect this tropifexor-induced, cell cycle-related response in models of liver injury in vivo (AMLN, [45]) and ex vivo (PCLS) (Supp. Table 5). BTEA of tropifexor-upregulated genes in both contexts rather reflected defense mechanisms against steroid, bile acid and xenobiotics accumulation. Why tropifexor, and hence FXR, loses its ability to induce cell cycle-related gene expression in injured tissues remains unclear. This may reflect an impaired release of priming growth factors by the liver or a context-dependent functional switch in FXR activity driven by mechanisms ranging from epigenomic remodeling to post-translational modifications. In line with this interpretation, the ability of tropifexor to upregulate fibrosis-related genes, including *Acta2, Col1a1, Col3a1*, and *Timp1*, in naïve livers also points to a functional switch in FXR activity between healthy and diseased tissues (Supp. Table 2), since tropifexor activity on these genes was clearly repressive in a fibrotic context. Consistent with this notion, we note that FXRα moderate overexpression in white adipose tissue depots stimulate extracellular matrix remodeling in chow-fed mice [94].

Liver fibrosis is driven by activation of hepatic stellate cells (HSCs), which convert from quiescent cells into extracellular matrix-producing myofibroblasts. Under chronic injury conditions, this wound-healing response becomes maladaptive and results in excessive extracellular matrix deposition and progressive alteration of liver structure and function. This transition is associated with metabolic reprograming and induction of a broad profibrotic transcriptional program, and translational strategies have therefore largely focused on targeting genes and pathways upregulated during HSC activation.

However, several pathways are attenuated during fibrosis progression, and we found that multiple intercellular communication (ICC) pathways are blunted in MASH and fibrosis while the FXRα isoform ratio is preserved. Among them, the chemerin pathway appeared consistently blunted in rodent models of MASH and fibrosis, a process likely explained by reduced expression of the preprochemerin-encoding gene *Rarres2*, which was also decreased in human steatotic and fibrotic livers. These findings are consistent with a previous report identifying *Rarres2* as a direct FXR target gene in HCs [95], which is indeed strongly inducible in in vivo tropifexor-exposed HCs and HSCs. Selective ablation of FXR^HC^ expression indicated that HSCs are a major site for *Rarres2* expression, and localization of chemerin receptors supports a HSC-Kupffer cells crosstalk in both human and mouse livers. The biology of chemerin in the liver remains incompletely understood, and the published literature is still contradictory [96]. One likely source of this variability is the complex intra- and extracellular processing of chemerin by multiple proteases, which generates isoforms with either agonistic or antagonistic activity [82]. The lack of a significant biological response to chemerin or the nonapeptide C9 in PCLS may reflect limited peptide stability, a constraint that was circumvented by using the CMKLR1 antagonist α-NETA. Although our results are consistent with tropifexor-induced preprochemerin synthesis followed by processing into a CMKLR1-antagonistic form, this interpretation remains to be formally demonstrated.

Taken together, our study expands the current understanding of FXR^HSC^ biology and suggests specific roles for bile acid nuclear signaling outside of the hepatocyte compartment, such as antioxidant responses and hepatoprotection. It also points to an important role of FXR^HSC^ in controlling hepatic ICC, whose integrity is essential for liver homeostasis.

## Acknowledgements

This work was supported by a grant from Fondation pour la Recherche Médicale (FRM, EQU202203014645) to PL, grants from Agence Nationale pour la Recherche (ANR “HSCReg”, ANR21-CE14-0032-01 to JE, ANR 10-LABX-0046 to BS), and grants from ERC [Advanced grant to BS (Immunobile n°694717), Starting grant to JTH (Metabo3DC n°101042759)].

## Declaration of interest

The authors declare no competing interests.

## Notes

### Competing Interest Statement

The authors have declared no competing interest.

