## Supplemental Information for "Hepatic stellate cell FXR signaling regulates context-dependent functions in liver homeostasis and fibrosis"

**to**

### MATERIAL and METHODS

**Reagents:** Tropifexor was obtained from TargetMol (#T4379). 2-(α-naphthoyl) ethyltrimethylammonium iodide (α-NETA), obeticholic acid, and GW4064 were purchased from Sigma-Aldrich (#SML2731, #SML3096, and #G5172, respectively). Carbon tetrachloride ($CCl_4$) was also sourced from Sigma-Aldrich (#289116).

**Antibodies:** Primary antibodies used throughout this study included anti-PKM2 (#4053, Cell Signaling Technology, RRID:AB_1904096), anti-phospho-PKM2 (Tyr105) (#3827, Cell Signaling Technology, RRID:AB_1904076), anti-SHP (#PA5-116825, Thermo Fisher Scientific), anti-FXR (#PP-A9033A-00, Perseus Proteomics; #ab187735, Abcam; or #72105, Cell Signaling Technology), anti-αSMA (#ab124964, Abcam, RRID:AB_11129103), and anti-HSP90AA1 (#662802, BioLegend), functional grade anti-TGFβ 1,2,3 (1D11.16.8, eBioscience™ # 16-9243-85, Invitrogen).

**Animal experimentation:** All animal studies were compliant with EU specifications regarding the use of laboratory animals and have been approved by the Nord-Pas de Calais Ethical Committee CEEA75. Unless stated otherwise, 8-12-week-old C57Bl6/J male mice from Charles River Laboratories were used for experimentation. Mice had either free access to water and to a standard chow diet (Safe Diet A04) (ad libitum conditions) or access to food was restricted to the active period for 2 weeks before euthanasia (12 h from ZT12 to ZT24) [time-restricted feeding (TRF) conditions], ZT0 being lights-on. Unless mentioned otherwise, animals were euthanized at ZT4-6 by cervical dislocation. Tissues were removed and snap-frozen in liquid nitrogen. Where indicated, wild type C57Bl6/J mice were fed either a control diet (standard rodent chow, 5% kcal fat) or a 'MASH' diet (45% kcal fat, 40% kcal carbohydrate, 15% kcal protein with 1% (by weight) cholesterol; SAFE Diets, France) for 24 weeks [1].

***Liver-specific FXR KO mice:*** Liver-specific FXR knockout mice and their control littermates were generated by cross-breeding mice harboring loxP sites flanking the final FXR exon ($FXR^{flox/flox}$) [2] with mice expressing Cre recombinase under the control of the albumin promoter [C57BL/6-Tg(Alb-Cre)21Mgn/J; Charles River, Portage, Michigan](Alb-$CRE^{+/-}$ mice) as described [3, 4].

***Intestinal-specific FXR KO mice:*** Intestinal-specific FXR knockout mice and their control littermates were generated by cross-breeding mice harboring loxP sites flanking the final FXR exon ($FXR^{flox/flox}$) [2] with mice expressing Cre recombinase under the control of the villin promoter [5].

***Tropifexor administration:*** An initial 1 mg/mL tropifexor stock solution was made by dissolving tropifexor in DMSO. This solution was diluted 8-fold in 45% propylene glycol (MP Biomedicals, #151957),

5% Tween 20 (Sigma, #1379) before injection. C57Bl/6J mice (12-week-old, n=6/group) were treated daily at ZT4 with 0.5 mpk tropifexor for 3 days. Livers were collected at ZT4-5 and either processed for hepatocytes and hepatic stellate cell purification or snap-frozen.

*$CCl_4$ administration:* A $CCl_4$ solution was prepared in filtered (0.22µm) olive oil (Sigma, #O1514)(1:8, vol:vol) and injected intraperitoneally (0.5mL/kg, 800 mpk) once or 3 times a week for 2 weeks. Livers were then processed for cell sorting and purification as described in [6].

**Mouse hepatocyte and hepatic stellate cell isolation:** Primary mouse HSCs were isolated from C57BL/6 J mice (male, 15–18-week-old, Charles River Laboratories) according to [6]. After cervical dislocation, livers were perfused through the vena cava first with HBSS (Gibco-ThermoFisher, #14170), then with a dissociation buffer [1x HBSS, pH 7.4, 1 mM $CaCl_2$, 100 U/mL collagenase type IV (Sigma #C5138)]at 37°C. Hepatocytes were pelleted after tissue mincing after a brief centrifugation (50xg for 2 min.) and resuspended first in red blood cell lysis buffer (155 mM $NH_4Cl$, pH 7.4, 10 mM $NaHCO_3$, 0.127 mM EDTA) then in fluorescence-activated cell sorting (FACS) buffer [1× phosphate-buffered saline (PBS), pH 7.4, 0.5% bovine serum albumin] supplemented with RNAsin (1:1000, Promega, #N2511). HSCs were separated from other non-parenchymal cells by FACS using an Aria II SORP (BD Biosciences, Franklin Lakes, NJ, USA) based on retinol autofluorescence by excitation with a 355 nm laser and detection with a 450/50 nm band-pass filter. Cell viability was estimated by 0.5% Zombie Green (BioLegend®) staining. A detailed procedure is available elsewhere [6].

**The human ABOS cohort:** The Hôpital Universitaire de Lille (HUL) cohort (ABOS; ClinicalTrials.gov: NCT 01129297) has been described in detail in [7-9]. Briefly, liver needle biopsies were obtained at the time of surgery from 910 obese patients undergoing bariatric surgery and processed for transcriptome analysis was performed using Affymetrix Human Transcriptome Array (HTA) 2.0. Livers were stratified using stringent qualitative and quantitative criteria into “healthy, normal”, “steatotic,” and “MASH” livers. The fibrosis stage was assessed using the Kleiner scoring system and was used to define a propensity score–matched subcohort based on age, sex, body mass index (BMI), diabetic status and statin use (see [10]). Fifty-three patients with MASH and fibrosis (F>2) were thus matched with 53 MASH-only patients (F=0) [11].

**RNAscope:** RNAscope assays on paraffin-embedded liver sections were performed using the RNAscope®Multiplex Fluorescent Reagent Kit v2 and probes according to the manufacturer’s instructions (Adv. Cell Diag., Bio-Techne). RNAscope® probes used were Hs-COL1A1-C2 (#401891-C2) and Hs-NR1H4-C1 (#494541). The anti-αSMA antibody used for immunofluorescence was from Abcam (ab124964 [EPR53681]) and used at 1:50 final dilution. Briefly, paraffin-embedded tissue sections were deparaffinized and treated with hydrogen peroxide to block endogenous peroxidase activity. Antigen retrieval was performed by incubating sections in RNAscope® 1× Target Retrieval Reagent at 99 °C for 15 minutes, followed by treatment with RNAscope® Protease Plus at 40 °C for 30 minutes in the HybEZ™ oven. Subsequently, tissue sections were hybridized with RNAscope® target probes according to the manufacturer's instructions. After hybridization, the signal was amplified through three sequential amplification steps. For detection, RNAscope® HRP-C1 was applied first, followed by incubation with Opal™ 650 fluorophore (Akoya Biosciences) for 30 minutes. After washing, HRP-C2 was applied, followed by incubation with Opal™ 570. Following RNAscope detection, sections were blocked in 10% normal goat serum overnight at room temperature. Sections were then incubated overnight at 4 °C with the anti-αSMA primary antibody (ab124964, clone EPR5368, Abcam) at 1:50 dilution. After washing, sections were incubated with a goat anti-rabbit Alexa Fluor® 488-conjugated secondary antibody at 1:1000 dilution for 1 hour at room temperature in the dark. Sections were mounted in antifade mounting medium (Vectashield® Antifade Mounting Medium, #H1000) and visualized with an Axioscan Z1 slide scanner (Zeiss).

**Cell lines, plasmids and transfection:**

*Cell lines:*

The mouse hepatic stellate cell line was obtained from Kerafast (#EMS404). Thereafter termed EMS404, this cell line isolated from *Tlr4*$^{-/-}$C57Bl6 mouse liver stably expresses human TLR4 and was immortalized using a pCMV-SV40 Large T antigen vector [12]. EMS404 cells were cultured in Dulbecco's Modified Eagle Medium (DMEM) supplemented with 5% fetal bovine serum (Dutscher). Mouse AML12 hepatocytes (RRID:CVCL_0140) were derived from normal hepatocytes of a transgenic C57BL/6 mouse that expressed the human transforming growth factor alpha (TGFα) gene [13]. They were maintained and expanded as previously described [14]. Where indicated, cells were synchronized by treatment with dexamethasone (100 nM) for 2 h prior to experimentation.

*Transfection:* EMS404 cells were cultured at 37 °C in 1x DMEM supplemented with GlutaMAX™ (Gibco™, #31966-021) supplemented with 5% fetal calf serum (FCS) (Dutscher™, #S1810-500), and 1% penicillin-streptomycin (Gibco™, #15140-122). For transfection, Lipofectamine® 2000 (Invitrogen™, #11668019) was used with plasmid DNA at a reagent-to-DNA ratio of 3:1. The reagents and DNA were diluted in Opti-MEM® Reduced Serum Medium (Gibco™, #31985-070), and cells were cultured for 18 hours in antibiotic-free complete medium. Twenty-four hours post-transfection, cells were treated for 24 hours with either GW4064 or tropifexor. Both compounds were dissolved in DMSO (Sigma-Aldrich®, #D8418) and diluted 1:1000 in the regular culture medium supplemented with 0.2% Bovine Serum Albumin (BSA) (Sigma-

Aldrich®, #A7030) in place of FCS. TGFβ treatment (stock solution: 10 µg/mL in vehicle: 4 mM HCl, 1 mg/mL BSA; R&D Systems®, #240-B-002) was performed on cells that had been serum-starved for 9 hours in DMEM supplemented with 1% penicillin-streptomycin and 0.2% BSA. Cells were then treated with TGFβ at a final concentration of 1 ng/mL for 24 hours in the same medium further supplemented with 0.5% fetal calf serum (FCS). Plasmids used in this study were: pcDNA3-FXRα1, pcDNA3-FXRα2, pcDNA3-FXRα3, pcDNA3-FXRα4 and pcDNA3-RXRα, all encoding for mouse *Nr1h4* transcripts. They have been described in [14, 15].

**RNA extraction and RT-qPCR:** RNA was extracted from cultured cells using the NucleoSpin® RNA Kit (Macherey-Nagel™, #740955) and from frozen mouse liver samples using TRIzol® Reagent (Ambion®, Thermo Fisher Scientific, #15596026). For the latter, each liver piece was homogenized using a T10 basic ULTRA-TURRAX® homogenizer (IKA®, #0003737000) in 600 µL of TRIzol®, followed by incubation for 5 minutes at room temperature. Subsequently, 120 µL of chloroform (Sigma-Aldrich, #C2432) was added, and the samples were centrifuged at 15,700 × g for 15 minutes at 4 °C. The upper aqueous phase was carefully recovered, and an equal volume of isopropanol (Sigma-Aldrich, #I9516) was added to precipitate the RNA. After incubation for 10 minutes at room temperature, samples were centrifuged again (15,700 × g, 15 minutes, 4 °C). The RNA pellet was washed twice with 70% ethanol (VWR, #20821.330), with centrifugation steps (15,700 × g, 5 minutes, 4 °C) between washes. Finally, RNA was resuspended in 100 µL of nuclease-free water (Thermo Fisher Scientific, #10977015). RNA quality was assessed by measuring the A260/A280 ratio (to detect protein contamination) and the A260/A230 ratio (to detect polysaccharide or phenol contamination). Reverse transcription (RT) was performed using the High-Capacity cDNA Reverse Transcription Kit (Applied Biosystems®, Thermo Fisher Scientific, #4368814), according to the manufacturer's instructions. Quantitative real-time PCR (qPCR) was carried out using PowerTrack™ SYBR™ Green Master Mix (Applied Biosystems, #A46109) on a QuantStudio™ 3 Real-Time PCR System (Applied Biosystems®, Thermo Fisher Scientific, #A28567). The expression levels of target genes were normalized to *Rps28* (40S ribosomal protein S28) or RPLP0 (Ribosomal Protein Lateral Stalk Subunit P0) as the reference genes. Relative gene expression was calculated using the $2^{-\Delta\Delta Ct}$ method [16].

**Transcriptomic analysis:**

***Affymetrix microarrays:*** GeneChip® Whole Transcript (WT) Expression Arrays (Affymetrix®-Thermo Fisher Scientific, #902281) were used in combination with the GeneChip® WT PLUS Reagent Kit (Affymetrix®, Thermo Fisher Scientific, #902280), following the manufacturer's protocol. A detailed procedure is available in [17]. Data processing was performed using our local GIANT analysis workflow [18]

*RNA sequencing:* Following initial quality control ($OD_{260/280}$ and RNA Integrity Number [RIN]), mouse RNA samples underwent oligo(dT) selection, random hexamer (N6) primed reverse transcription, end repair, and adapter ligation. Libraries were sequenced using paired-end 100 bp reads on the DNBSEQ platform (BGI, Shenzhen, China). Sequence data were quality-filtered using SOAPnuke (v1.5.2) with the parameters: -l 15 -q 0.2 -n 0.05. Cleaned reads were stored in FASTQ format. Read alignment to the mouse reference genome (mm10) was performed using HISAT2 (v2.0.4)[19] with the following parameters: --sensitive --no-discordant --no-mixed -I 1 -X 1000 -p 8 --rna-strandness RF. For transcriptome mapping, Bowtie2 (v2.2.5) was used with the parameters: -q --phred64 --sensitive --dpad 0 --gbar 99999999 --mp 1,1 -np 1 --score-min L,0,-0.1 -p 16 -k 200. Gene expression levels were quantified using RSEM (v1.2.8) [20]. Sample collection and data processing for the RNA-seq analysis of human liver biopsies have been described elsewhere [7]. FXR isoform expression levels were estimated using the RSEM v1.3.0 “rsem-calculate-expression” function for paired end sequencing (--paired-end --strandedness RF) with alignments mapped to the human genome (GRCh38) with STAR v2.7.3a and quantified using the Ensembl 105 transcript annotation database.

**Protein extraction, western blotting and WES analysis:**

***Total protein extraction:*** Cells were washed twice with 1x DPBS (Thermo Fisher Scientific™, #14190144) and lysed for 30 minutes on ice using the following lysis buffer: 100 mM Tris-HCl, pH 7.4, 100 mM NaCl, 2 mM EDTA (pH 8.0), 25 mM NaF, 0.1% Triton X-100, 1 mM benzamidine, 0.1 mM $Na_3VO_4$, and 1x protease inhibitor cocktail (Roche, #11836153001). Lysates were centrifuged at 16,000 × g for 20 minutes at 4 °C to remove cellular debris.

*Protein extraction with cell fractionation:* Cells were collected in 1x DPBS, centrifuged at 600 × g for 2 minutes at 4 °C, and resuspended in Buffer A (10 mM HEPES, pH 7.9, 10 mM KCl, 1.5 mM $MgCl_2$, 0.34 M sucrose, 10% glycerol, 1 mM DTT, 1X protease inhibitors, 0.1% Triton X-100). After incubation on ice for 10 minutes, samples were centrifuged at 1,300 × g for 5 minutes at 4 °C. The supernatant (cytoplasmic fraction) was recovered, while the nuclear pellet was washed again with Buffer A. The cytoplasmic fraction was cleared of organelles by further centrifugation at 16,000 × g for 5 minutes at 4 °C. The nuclear pellet was incubated for 30 minutes on ice in Buffer B (10 mM HEPES, pH 8.0, 3 mM EDTA, 0.2 mM EGTA, 1 mM DTT, and protease inhibitors). After centrifugation at 1,700 × g for 5 minutes at 4 °C, the supernatant (nucleoplasm) was collected, and the chromatin pellet was washed once with Buffer B and once with cold 1x DPBS. The pellet was then resuspended in digestion buffer [50 mM Tris-HCl, pH 8.0, 1 mM $MgCl_2$, and benzonase (2.75 U/µL; Sigma-Aldrich, #E1014] for 20 minutes at room temperature to digest nucleic acids. The reaction was stopped by adding Laemmli 6X buffer (Bio-Rad, #1610747) before SDS-PAGE separation.

*Western Blot:* Protein concentration was determined using the Pierce™ BCA Protein Assay Kit (Thermo Fisher Scientific™, #23225), using BSA as a standard. Proteins were resolved by SDS-PAGE and transferred onto nitrocellulose or PVDF membranes using the iBlot® 2 Transfer System and iBlot® 2 NC or PVDF Mini/Regular Stacks (Invitrogen™, #IB23001 or IB24001). Membranes were incubated with primary antibodies as indicated, diluted in 5% BSA (Sigma-Aldrich, #A7906) in 0.1% TBST (Tris-buffered saline with 0.1% Tween-20). Detection was performed by enhanced chemiluminescence using one of the following Thermo Fisher Scientific™ reagents, depending on the sensitivity required: Pierce™ ECL Western Blotting Substrate (#32106), or SuperSignal™ detection systems (SuperSignal™ West Femto, #34096 or West Atto, #A38555). Images were acquired using the iBright™ FL1500 Imaging System (Invitrogen™, #A44115).

*Wes detection system:* For primary cell protein extracts, quantification was performed using the Pierce™ Micro BCA™ Protein Assay Kit (Thermo Fisher Scientific™, #23235) with BSA as a standard. Protein analysis was performed on the Wes™ system (ProteinSimple®, Bio-Techne), following the manufacturer's protocol.

**Preparation and culture of precision-cut liver slices:**

Male C57BL/6J mice were sacrificed by cervical dislocation at Zeitgeber Time (ZT) 5–7, and livers were immediately harvested and placed in ice-cold Hanks' Balanced Salt Solution (HBSS). One liver lobe per mouse was sectioned into 200 µm-thick slices using a vibratome tissue slicer (Campden Instruments, London, UK) under the following settings: 80 Hz frequency, 0.33 mm/s speed, and 4 °C. Each slice was then transferred onto a Millicell® cell culture insert (Millipore, #PICMORG50) placed in a 35 mm culture dish containing Dulbecco's Modified Eagle Medium (DMEM) (Sigma-Aldrich, #D5030), supplemented with 10 mM HEPES, pH 7.4, 25 mM glucose, 10% fetal calf serum, 4 mM sodium bicarbonate, 1% Glutamax and 1% penicillin-streptomycin. For viability assessment, pilot experiments were first performed using livers from Per2::Luc reporter mice [21]. In these cases, the culture medium was further supplemented with 200 µM beetle luciferin potassium salt. PCLS were cultured at 36°C without $CO_2$ for several days in a regular incubator or in a KRONOS-DIO AB-2550 system (Atto)(see [22] for details).

**Statistical analysis:**

Unless otherwise stated, statistical analyses were performed using GraphPad Prism (version 9 or later). Data are presented as mean ± standard error of the mean (SEM). At least three independent experimental replicates were performed. For in vitro data, equality of variances was assessed using the F test. Two-group comparisons were performed using an unpaired two-tailed t-test with Welch's correction. A one-way ANOVA followed by Tukey's multiple comparisons test (comparing all groups) was used for multiple comparisons involving one variable. When more than one variable was analyzed, a two-way ANOVA followed by Tukey's multiple comparisons test was applied.

**Data analysis:**

*Differential gene expression analysis:* After sequencing, raw data files were processed as described above to generate gene-level count matrices. These count files were uploaded to the Omics Playground v. 3.0 (BigOmics Analytics) [23] for downstream analyses and generation of visual representations. VolcaNoseR was used to generate and label Volcano plots [24].

*NicheNet*: NicheNet [25] was used with minor adaptations. The original ligand-target matrix was replaced with a modified, hand-curated version of the CellTalk database, restricted to mouse-specific ligand–receptor pairs [26]. The script was further adapted for bulk tissue analysis, where the same cellular population acts as both "sender" and "receiver". Circos plots and other data visualizations were generated using NicheNet and edited in CorelDRAW to improve readability.

*Isceberg*: Interactive visualization and exploration of single-cell RNA-seq datasets were performed using the ISCEBERG (Interactive Single Cell Expression Browser) R/Shiny application [27]. Raw or preprocessed Seurat objects were uploaded to the platform, which enabled quality control, clustering, dimensionality reduction (UMAP/t-SNE), marker gene identification, and differential expression analysis. Annotation was performed manually or using integrated tools (e.g., SCINA). Plots and results were exported for downstream analysis and figure preparation.

**Datasets:**

Publicly available datasets used in this study included: the transcriptomic analysis of liver FXR knockout mice [4] (EMBL-EBI: E-MTAB-1722), of mouse livers from the amylin NASH model [28] (NCBI-GEO: GSE129389), of mouse livers under time-restricted feeding [6] (NCBI-GEO: GSE223360), of single-cell and single-nucleus from mouse and human livers [29] (NCBI-GEO: GSE192742) and FXR ChIP-seq analysis in control and TCA-treated mouse livers [15] (NCBI-GEO: GSE87866).

Datasets generated as part of the present study included microarray analysis of total mRNA from EMS404 cells overexpressing FXRα1 or FXRα2 and treated or not with GW4064 (NCBI GEO: GSE331297); RNA-seq analysis of total mRNA from vehicle- or tropifexor-treated precision-cut liver slices (PCLS) prepared from wild-type and hepatocyte-specific FXR knockout mice (NCBI GEO: GSE333328); RNA-seq analysis of liver mRNA from wild-type and hepatocyte-specific FXR knockout mice (NCBI GEO: GSE332832); RNA-seq analysis of hepatocyte mRNAs isolated from vehicle- and tropifexor-treated wild-type mice (NCBI GEO: GSE334504); RNA-seq analysis of hepatic stellate cells isolated from vehicle- or tropifexor-treated mice (NCBI GEO:

GSE334831), RNA-seq analysis of liver mRNA from wild-type mice treated with vehicle or tropifexor (GEO NCBI: GSE335123), RNA-seq analysis of liver mRNA from wild-type mice treated with vehicle or OCA (GEO NCBI: GSE335333).

### SUPPLEMENTAL FIGURES AND TABLES

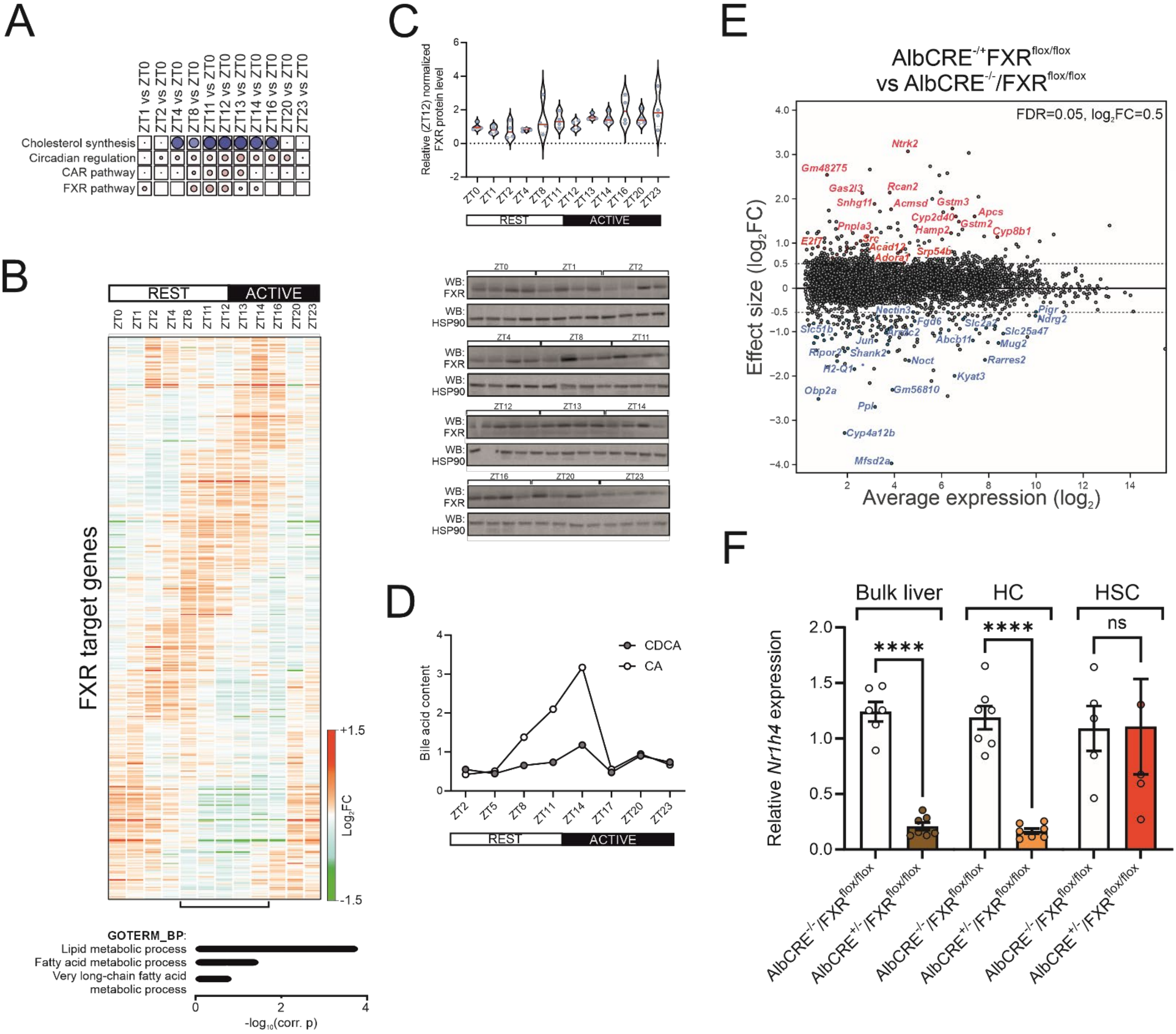

**Supplemental Figure 1: The FXR signaling pathway in mouse liver.** A) Time-of-day-dependent activation of relevant biological processes in mouse liver. The mouse liver transcriptome was analyzed using DNA microarrays at multiple time points (n=6 per condition) across a 24-hour period (ZT1, ZT2, ZT4, ZT8, ZT11, ZT12, ZT14, ZT16, ZT20, and ZT23), each compared with ZT0. Differentially expressed genes (|FC|>1.5, q value<0.05) were functionally annotated using the Gene Ontology Biological Process database and clustered accordingly. Blue, downregulated; red, upregulated. B) Heatmap of FXR target gene expression across the day-night cycle (upper panel). Upregulated genes at the fasting-feeding transition (ZT8-ZT14) were functionally annotated using the Gene Ontology Biological Process database, and the most significantly enriched terms are shown (lower panel). C) FXR protein expression across the day-night cycle (n=3 per condition). D) Principal primary bile acids content in mouse liver across the day-night cycle. E) MvA plot of gene expression in wild type (AlbCRE$^{-/-}$ FXR$^{flox/flox}$) versus FXR$^{HC}$ KO (AlbCRE$^{-/+}$ FXR$^{flox/flox}$) mice (n=3-6). Difference among means were compared using an unpaired two-tailed t-test (p<0.001: ****). Blue, downregulated; red, upregulated. F) *Nr1h4* transcript levels in bulk liver, purified HC and HSC from wild type and FXR$^{HC}$ KO mice.

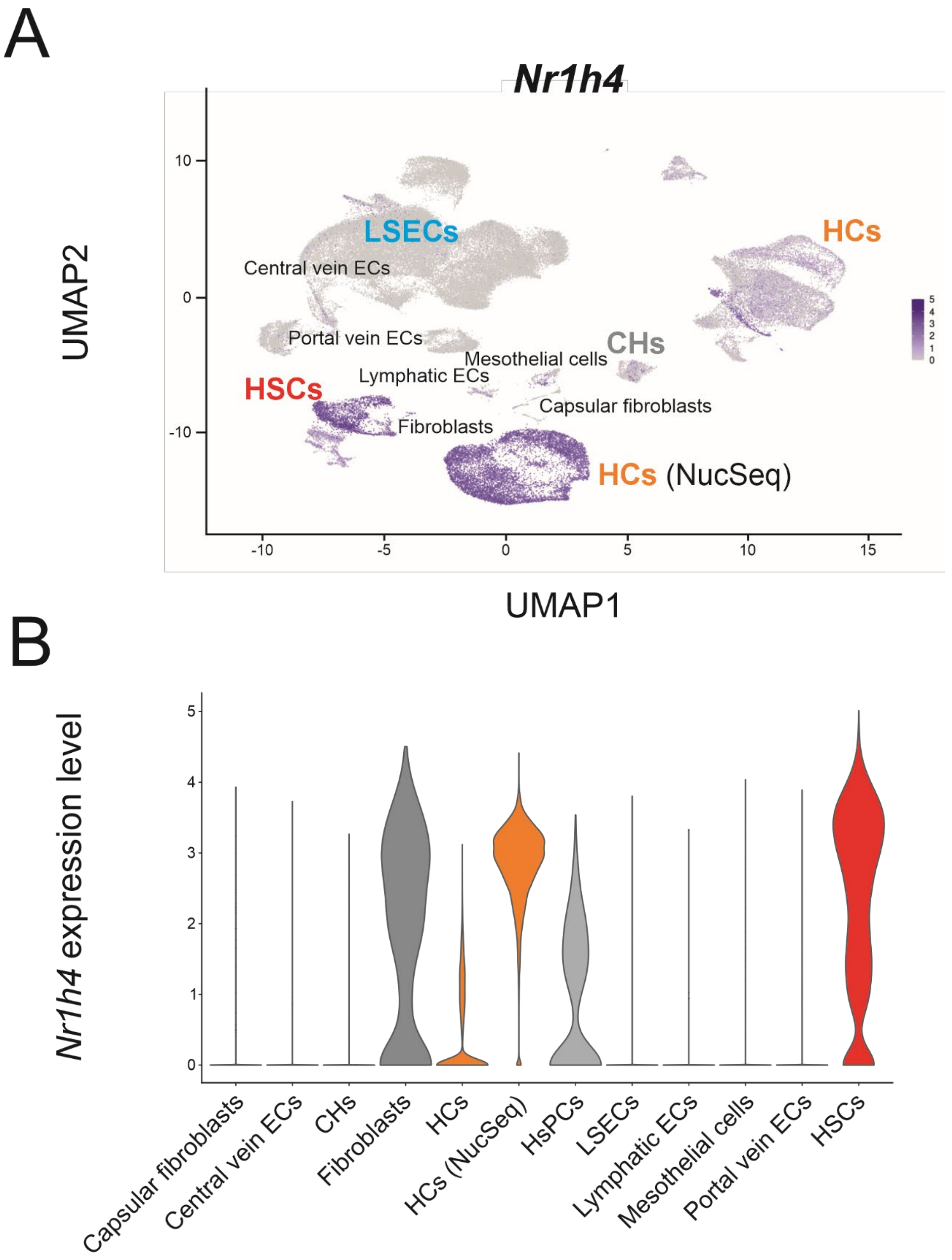

**Supplemental Figure 2: Single cell/single nucleus sequencing of mouse liver cells.** Sequencing data from mouse liver were extracted from the VIB database (https://www.livercellatlas.org) and reprocessed using our analysis pipeline, Isceberg. A) UMAP plot of *Nr1h4* expression across mouse liver cell types. B) Violin plot of *Nr1h4* expression level across mouse liver cell types.

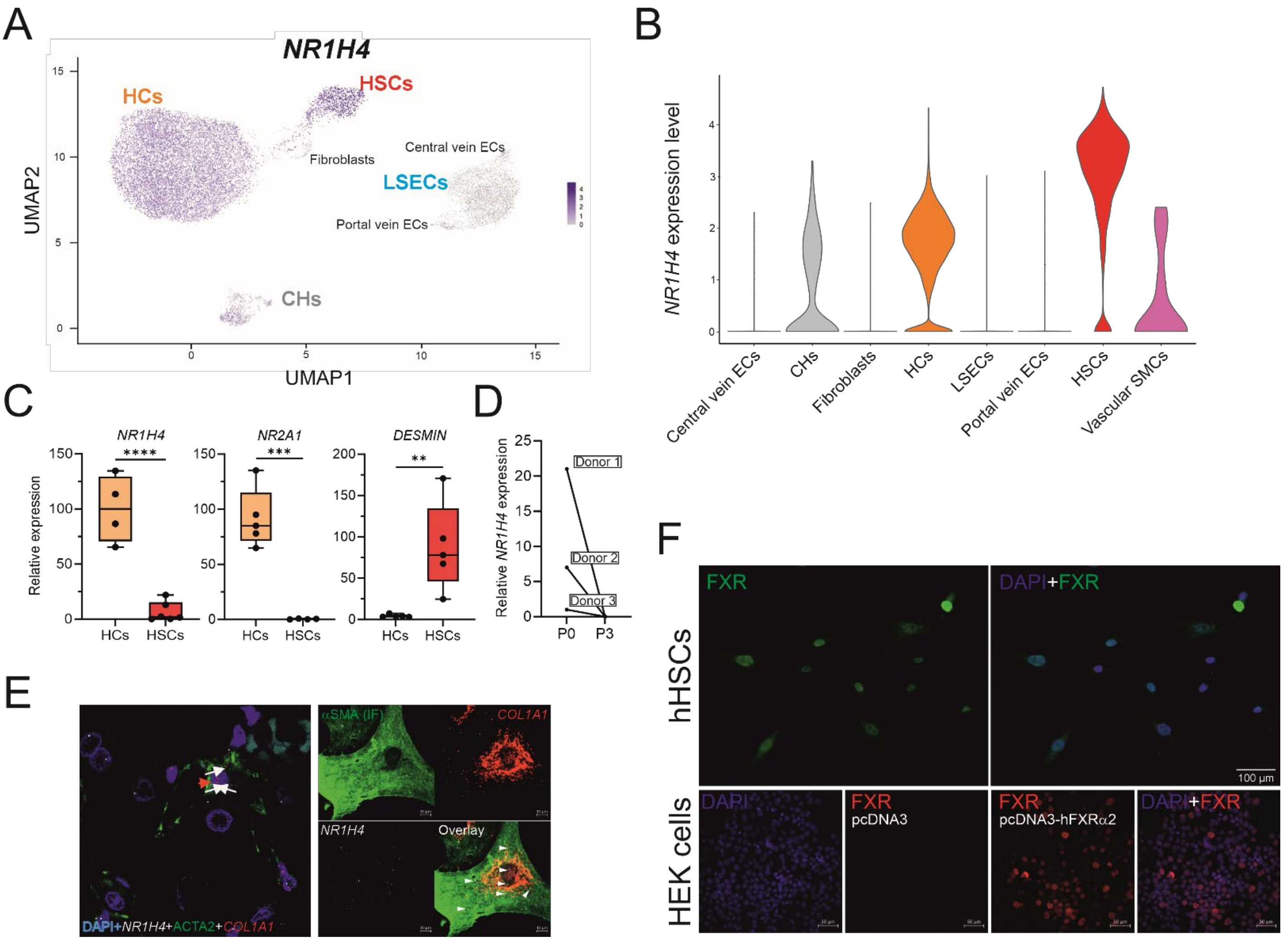

**Supplemental Figure 3: *NR1H4* expression in human liver cell types.** A) Sequencing data from human liver were extracted from the VIB database (https://www.livercellatlas.org) and reprocessed using our analysis pipeline, Isceberg. A) UMAP plot of *NR1H4* expression across human liver cell types. B) Violin plot of *NR1H4* expression level across human liver cell types. C) Gene expression in commercially available human HCs and HSCs. *NR1H4* (all isoforms), *NR2A1* (HNF4α, HC marker), *DESMIN* (HSC marker) expression levels were determined by RT-qPCR. Difference among means (n=4) was compared using an unpaired two-tailed t-test (p<0.01: **; p<0.005: ***; p<0.001: ****). D) *NR1H4* expression in various purified human HSC preparations. *NR1H4* expression was determined by RT-PCR at the lowest passage (P0) and after 3 passages. E) In situ hybridization in purified human HSCs. *NR1H4* and *COL1A* transcripts were localized using the RNAscope technology in donor 1 HSCs (P0). αSMA was localized by immunofluorescence. White arrowheads point at *NR1H4* transcripts. F) FXR protein expression in human HSCs. The FXR protein was detected by immunofluorescence in Donor 1 HSCs (P0) (upper panels). The specificity of the FXR antibody was validated in HEK cells transfected with either a FXR expression vector(pcDNA3-hFXRα2) or a control vector (pcDNA3).

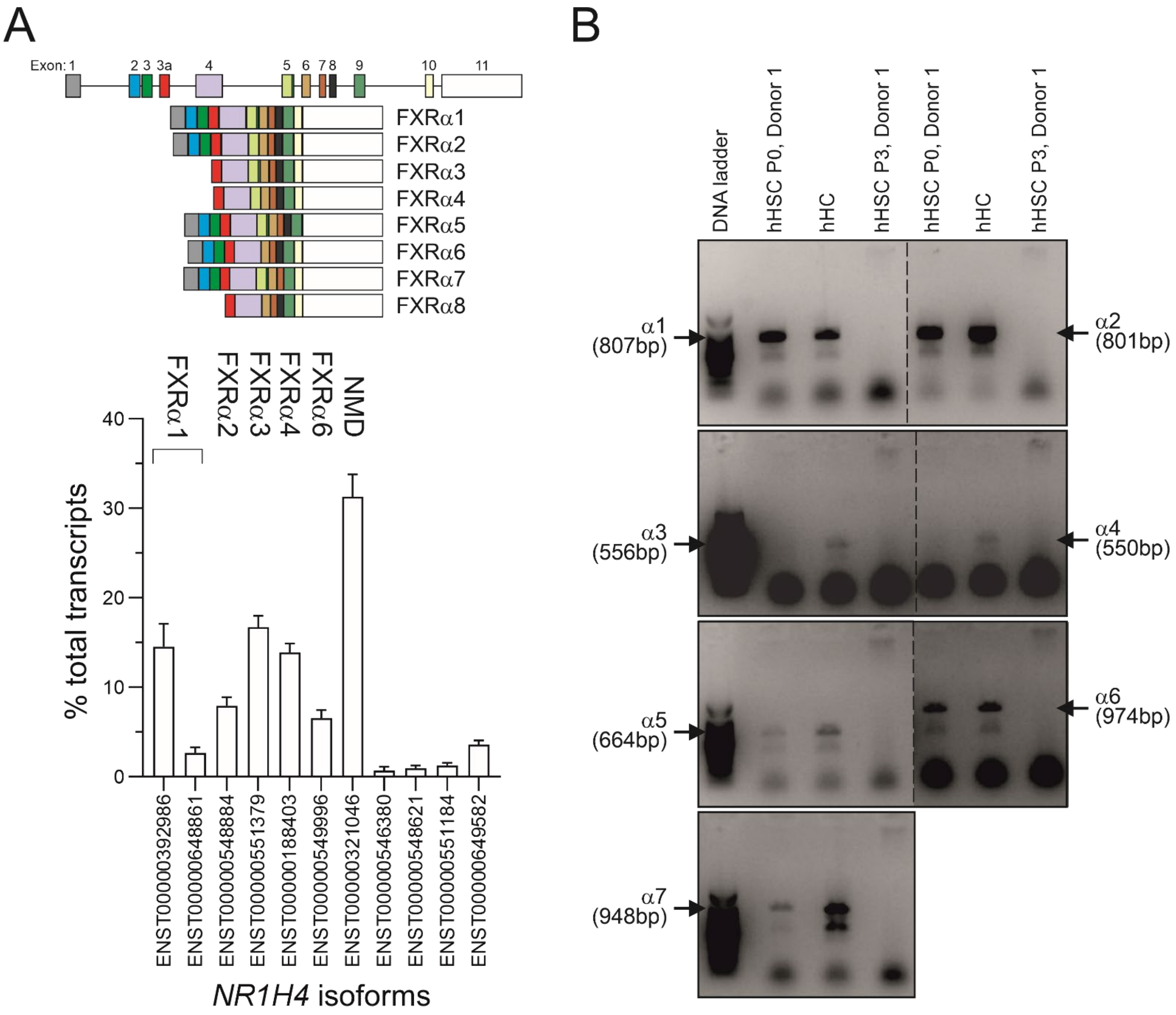

**Supplemental Figure 4: Expression of *NR1H4* isoforms in human liver and HSCs**. A) Schematic representation of *NR1H4* transcript isoforms (upper panel) (taken from [30]). *NR1H4* isoforms in human histologically normal livers. RNA-seq data were processed to identify NR1H4 isoforms. α5, α6 and α7 isoforms correspond to DBD or LBD-truncated mutants (lower panel). B) RT-PCR analysis of *NR1H4* isoforms in purified human HCs or HSCs. Primer sets were designed to amplify the indicated transcript.

| Gene Symbol | log2FC (GW4064 vs DMSO) | log2FC (α1_GW vs α1_DMSO) | log2FC (α2_GW vs α2_DMSO) | |
|---|---|---|---|---|
| *Gsta2* | 0,23 | 1,69 | -0,18 | FXRα1 |
| *Spink14* | 0,02 | 0,87 | 0,03 | |
| *Gstm1* | 0,08 | 0,84 | 0,18 | |
| *Olfr1166* | -0,31 | 0,81 | -0,11 | |
| *Myo15b* | -0,08 | 0,81 | 0,20 | |
| *Slco6d1* | -0,23 | 0,81 | 0,07 | |
| *Glipr1l3* | -0,23 | 0,76 | 0,25 | |
| *Tmem156* | -0,27 | 0,76 | -0,14 | |
| *Lzts3* | -0,19 | 0,74 | -0,04 | |
| *Cldn34c4* | -0,10 | 0,68 | -0,23 | |
| *Dhrs9* | -0,16 | 0,67 | -0,31 | |
| *Abcb1a* | 0,41 | 0,18 | 1,38 | FXRα2 |
| *Postn* | 0,93 | -0,08 | 1,21 | |
| *Disp2* | -0,13 | -0,06 | 1,20 | |
| *Map3k13* | 0,13 | 0,15 | 1,13 | |
| *Aqp1* | 0,14 | 0,06 | 1,05 | |
| *Acap1* | 0,18 | 0,23 | 1,05 | |
| *Asgr2* | 0,25 | 0,06 | 1,03 | |
| *Slco3a1* | -0,19 | 0,25 | 0,93 | |
| *Neurl3* | 0,16 | 0,05 | 0,88 | |
| *Nfkbiz* | 0,02 | 0,19 | 0,87 | |
| *Fabp6* | 0,62 | 5,58 | 9,09 | FXRα1+FXRα2 |
| *Osgin1* | 1,45 | 3,04 | 2,30 | |
| *Nppb* | -0,16 | 2,82 | 1,72 | |
| *Alpl* | 0,29 | 1,28 | 1,69 | |
| *Slc51b* | 0,32 | 1,07 | 3,03 | |
| *Fgfbp1* | -0,20 | 1,06 | 1,22 | |
| *Prl2c3* | -0,09 | 1,05 | 1,06 | |
| *Il4ra* | 0,32 | 0,99 | 1,56 | |
| *Slc51a* | 0,36 | 0,96 | 2,74 | |
| *Prl2c2* | -0,17 | 0,93 | 1,11 | |
| *Rbp4* | 0,69 | 0,91 | 2,87 | |

**Supplemental Figure 5: Selective gene activation by FXR isoforms.** EMS404 cells were transfected with the indicated FXRα expression vector and treated or not by GW4064 overnight. Transcripts were quantified by DNA microarrays and $log_2$(fold change, FC) were calculated after normalization (n=3, q value< 0.05). Genes were clusterized according to their isoform selectivity. Red: upregulated, green: downregulated.

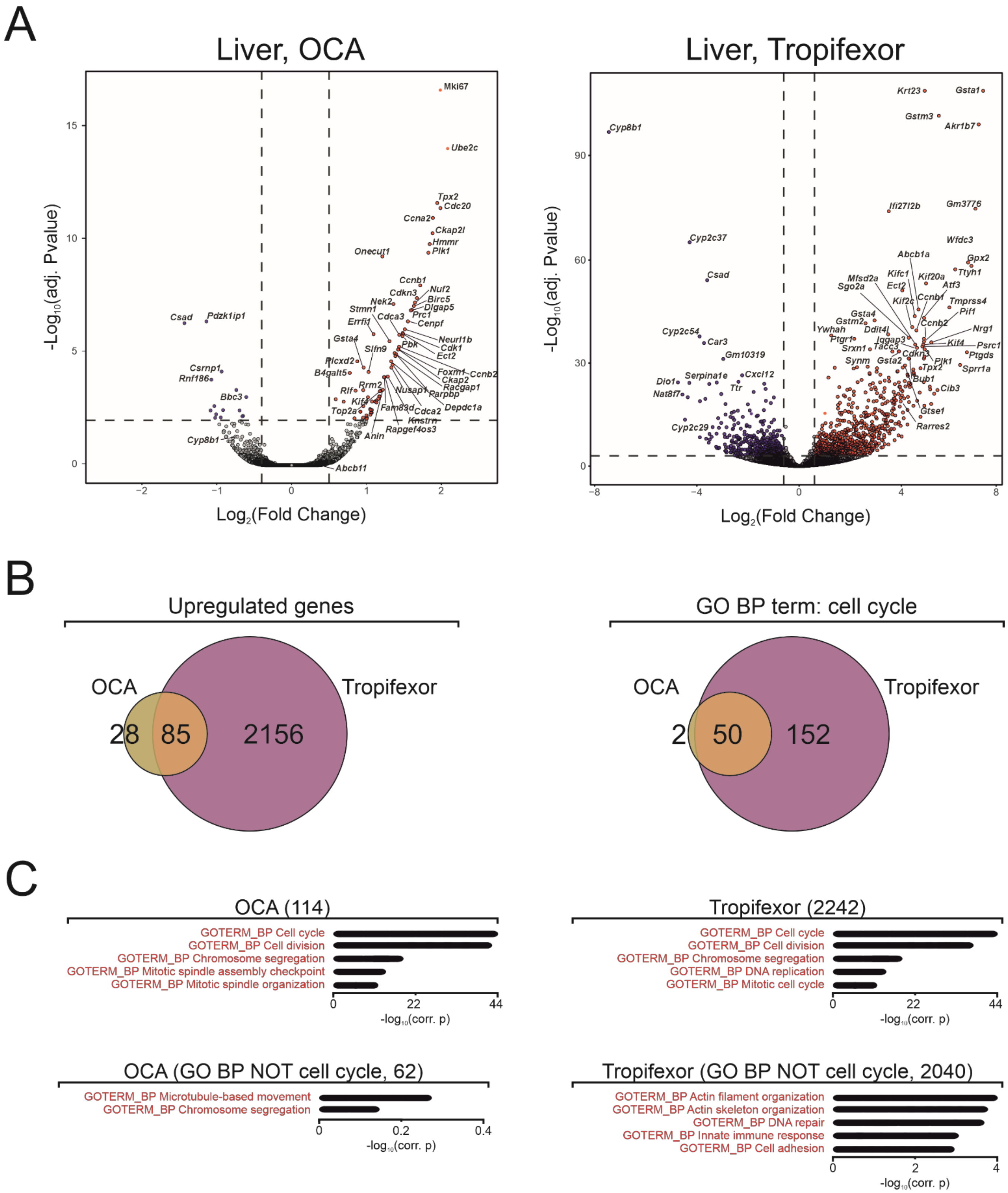

**Supplemental Figure 6: Tropifexor is a more potent FXR agonist in vivo than OCA**. A) Liver transcriptome after a short-term challenge with FXR agonists. Male mice were treated for 3 days with either OCA (30 mpk) or tropifexor (0.5 mpk). Livers were collected (n=4-6 per group) and the transcriptome was analyzed by bulk RNAseq. Differentially expressed genes are shown (FC>1.2, q<p0.05). Red: upregulated, blue downregulated. B) Venn diagram of upregulated genes after OCA or tropifexor treatment (left panel). Venn diagram of upregulated genes related to cell cycle control after OCA or tropifexor treatment (right panel). C) Biological term enrichment analysis (BTEA) of upregulated genes after OCA or tropifexor treatment. Upper panels: BTEA was performed using the GO Biological Process database without filtering. Lower panels: BTEA was performed after filtering out all genes related to cell cycle control.

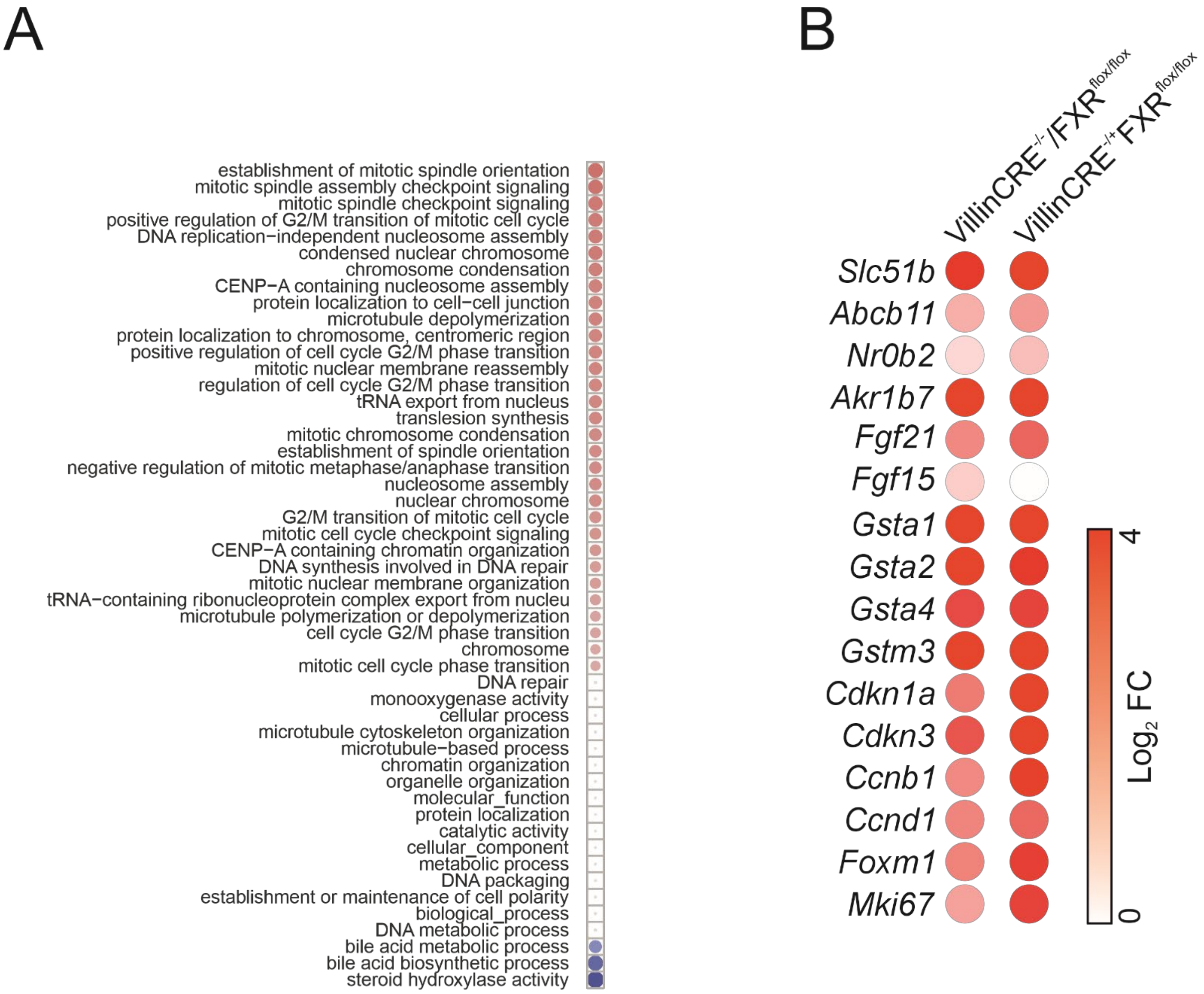

**Supplemental Figure 7: Tropifexor activity in wild-type and intestinal FXR knockout mice.** A) Activation matrix of hepatic biological pathways. GO terms consistently up- or downregulated in the livers of tropifexor-treated wild-type mice relative to vehicle-treated mice are shown (red, upregulated; blue, downregulated). (B) Hepatic gene expression in control and intestinal FXR knockout mice. Mice were treated with tropifexor for 3 days (0.5 mpk), and liver gene expression was analyzed by RT-qPCR. The bubble plot compares the fold changes measured in each genotype Red: upregulated.

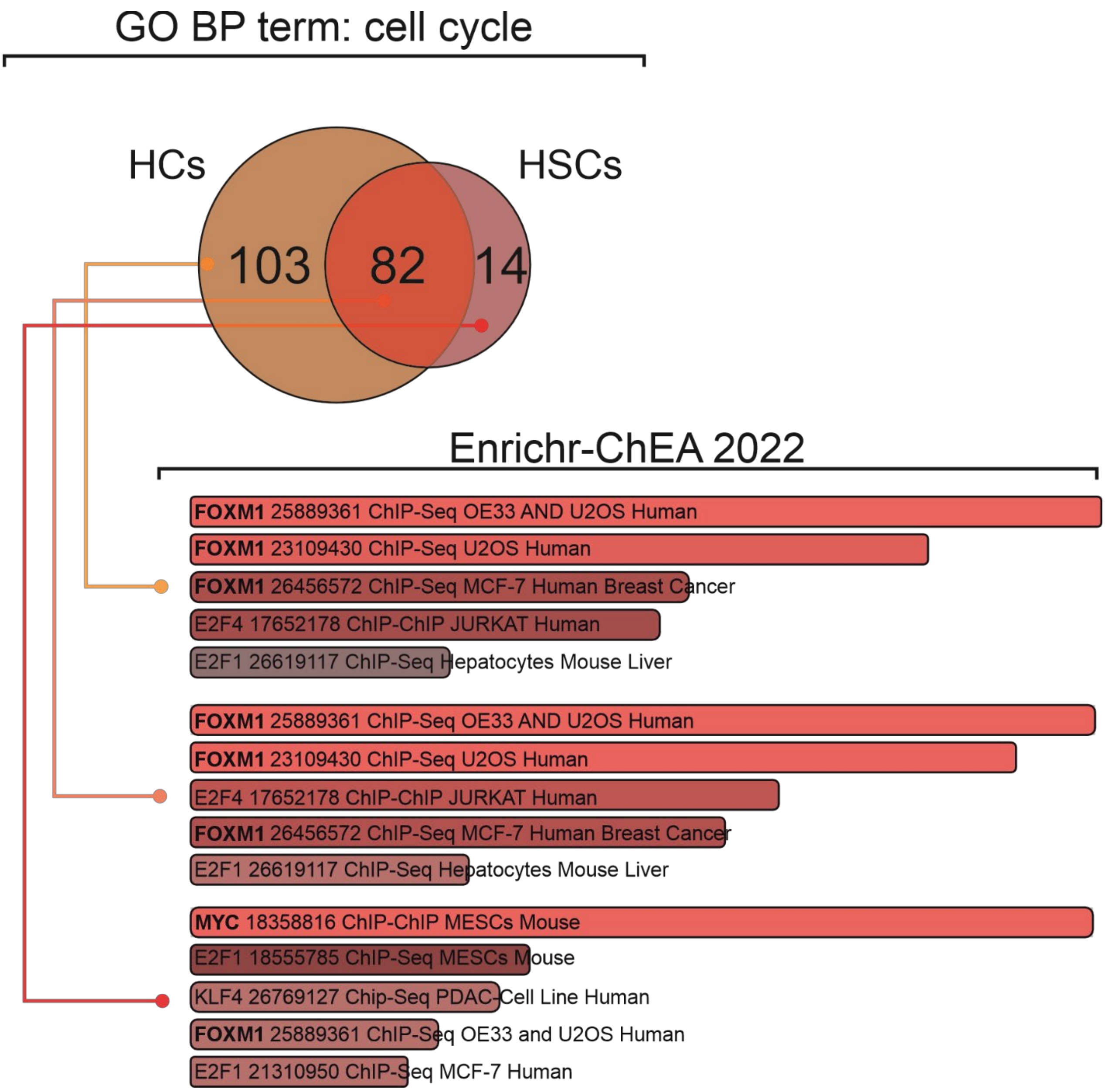

**Supplemental Figure 8: Putative drivers of the tropifexor-induced proliferative response.** Upregulated genes in HCs and/or HSCs related to cell cycle regulation were analyzed by ChIP-X enrichment analysis using Enrichr [31]. The most significantly overrepresented target gene sets associated with the indicated transcription factors are shown for HC-specific genes, HSC-specific genes, and commonly differentially expressed genes.

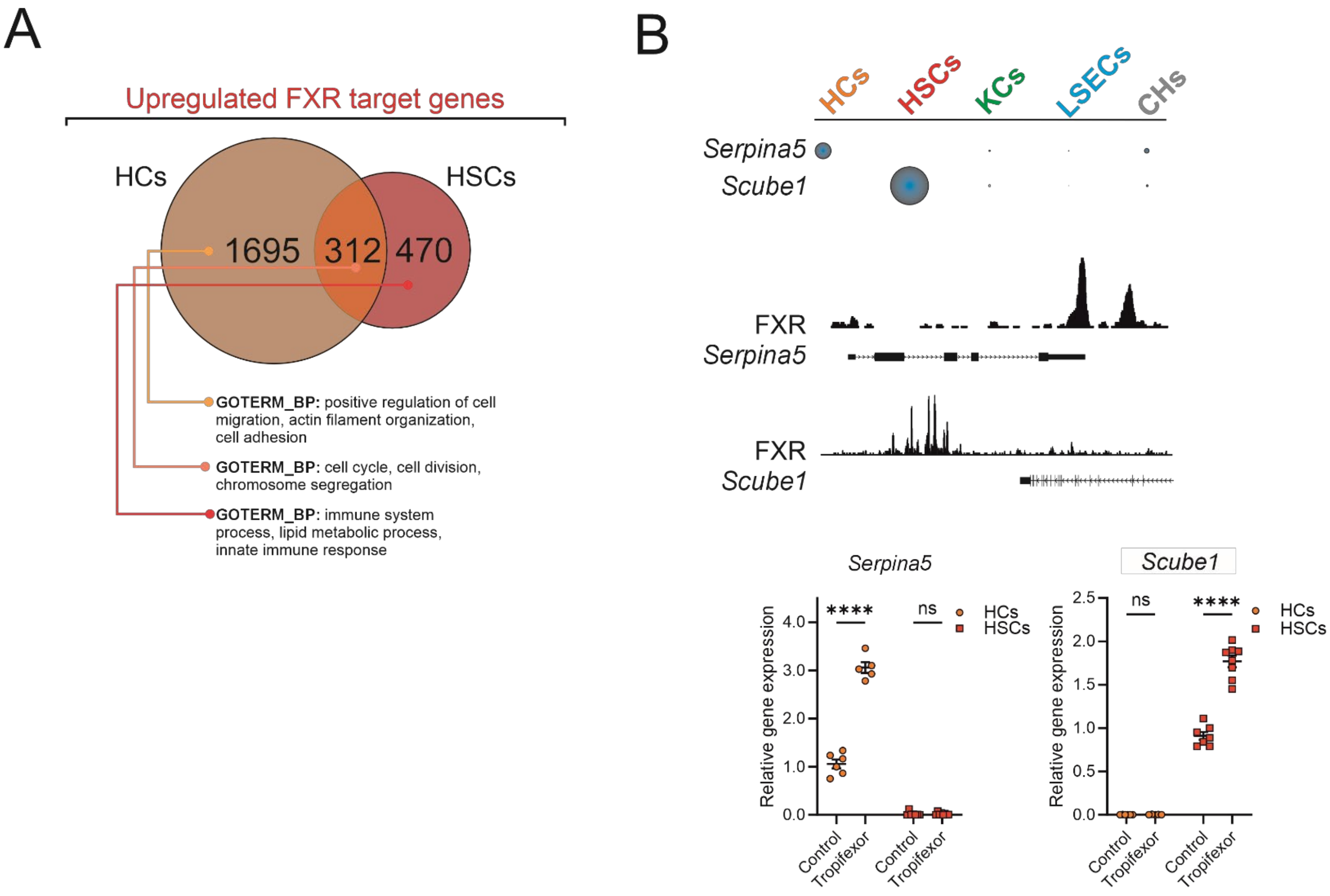

**Supplemental Figure 9**: Identifying HC- or HSC-specific FXR target genes. A) Refined BTEA analysis. Differentially expressed genes in HCs and HSCs following tropifexor treatment were intersected with a list of putative direct FXR target genes, defined by transcription start sites (TSSs) located near FXR ChIP-seq binding sites within 1 Mb in unchallenged mouse livers. BTEA was performed as in Figure 3C, and the top three hits from the Biological Process database are shown. B) Cell-specific expression of FXR-regulated genes. Basal expression levels of *Serpina5* and *Scube* 1 across mouse liver cell types are shown (upper panel), the corresponding whole liver FXR ChIP-seq profiles are shown in the middle panel, and their induction following tropifexor treatment in vivo is shown in the lower panel. Difference among means (n=6) were compared using a one-way ANOVA followed by a Tukey's multiple comparison test (p<0.001: ****).

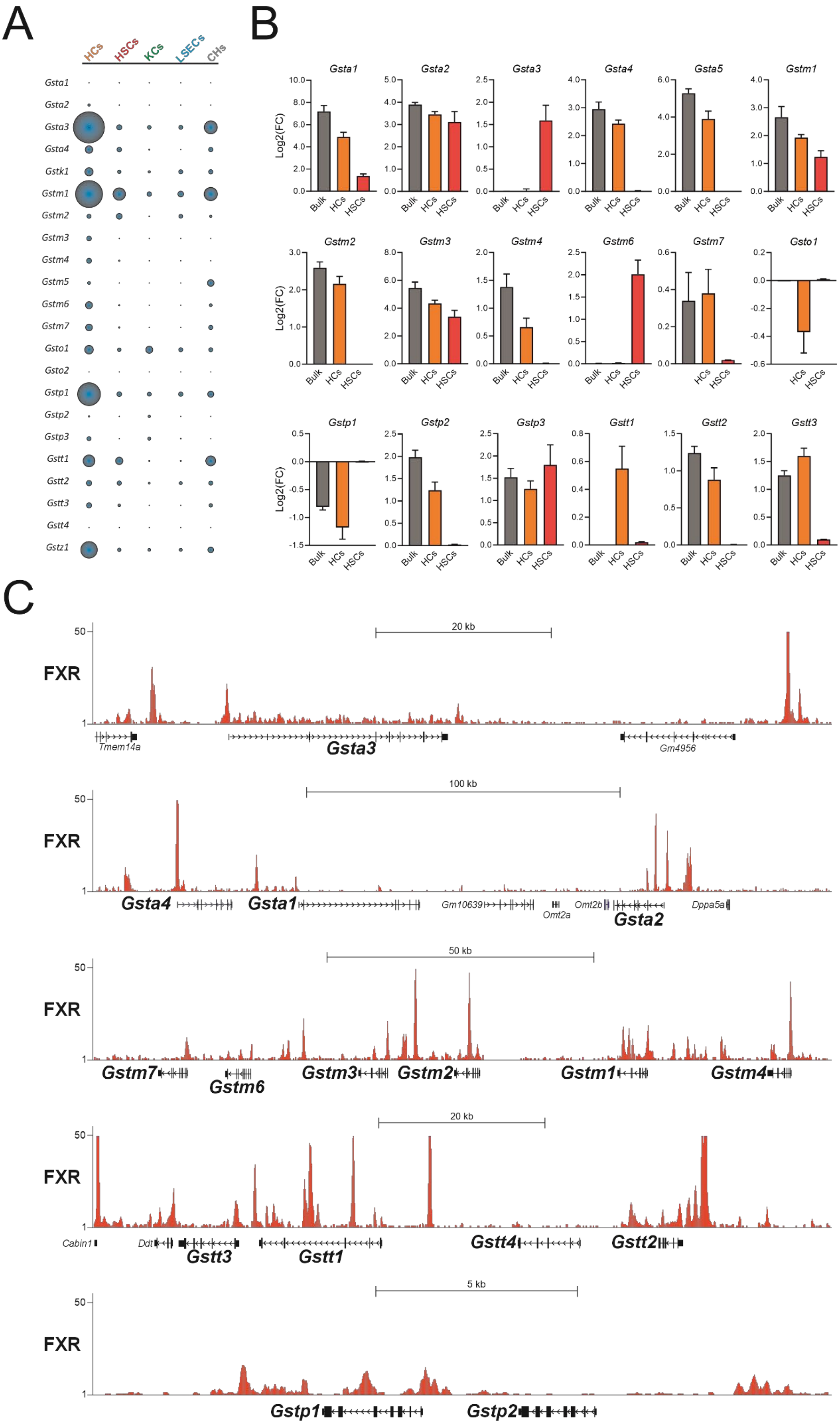

A
HCs
HSCs
KCs
LSECs
CHs
Gsta1
Gsta2
Gsta3
Gsta4
Gstk1
Gstm1
Gstm2
Gstm3
Gstm4
Gstm5
Gstm6
Gstm7
Gsto1
Gsto2
Gstp1
Gstp2
Gstp3
Gstt1
Gstt2
Gstt3
Gstt4
Gstz1
B
Log2(FC)
Gsta1
Gsta2
Gsta3
Gsta4
Gsta5
Gstm1
Gstm2
Gstm3
Gstm4
Gstm6
Gstm7
Gsto1
Gstp1
Gstp2
Gstp3
Gstt1
Gstt2
Gstt3
Bulk
HCs
HSCs
C
FXR
20 kb
Tmem14a
Gsta3
Gm4956
100 kb
Gsta4
Gsta1
Gm10639
Omt2a
Omt2b
Gsta2
Dppa5a
50 kb
Gstm7
Gstm6
Gstm3
Gstm2
Gstm1
Gstm4
20 kb
Cabin1
Ddt
Gstt3
Gstt1
Gstt4
Gstt2
5 kb
Gstp1
Gstp2

**Supplemental Figure 10: Cell-specific regulation of GST enzyme-encoding genes.** A) Cell-specific expression of GST isoforms across mouse liver cell types. B) Responsiveness of GST-encoding genes to tropifexor treatment in vivo. Observed $log_2$ fold changes in bulk liver tissue, isolated HCs, and isolated HSCs after in vivo treatment are shown. C) Whole-liver FXR ChIP-seq profiles at GST isoform loci in unchallenged livers.

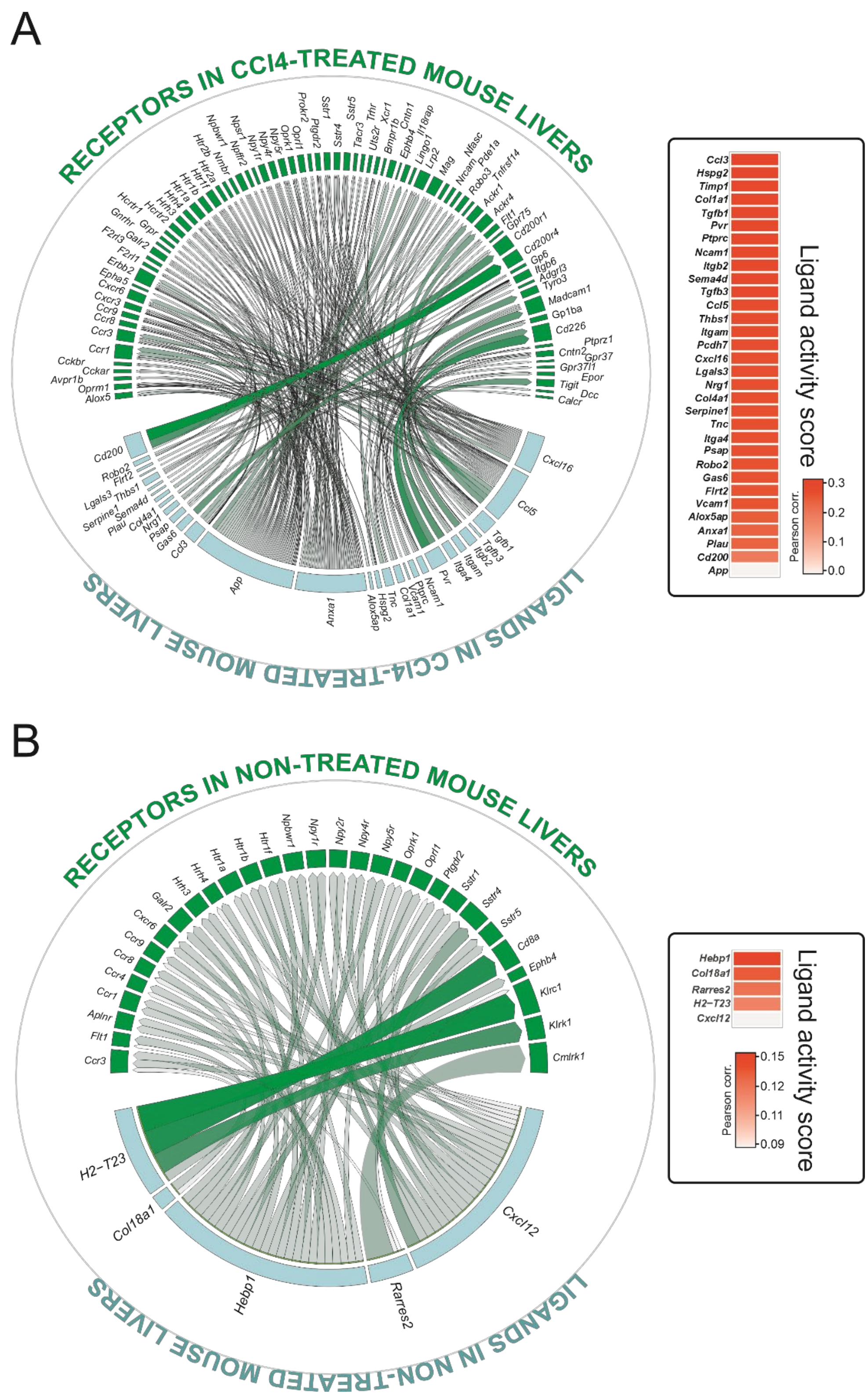

**Supplemental Figure 11: Ligand-receptor interactions in chemically induced fibrosis.** Gene expression in livers from vehicle- or $CCl_4$-treated male mice was assessed by RNA-seq and ligand-receptor (L-R) interactions were analyzed using the R package NicheNet adapted here for bulk tissue characterization. Relevant ligand-receptor pairs and associated ligand activity score were prioritized according to their predicted ability to account for the observed gene-expression changes. A) L-R interactions and ligand activity scores in $CCl_4$-exposed mouse livers. B) L-R interactions and ligand activity scores in vehicle-exposed (control) mouse livers.

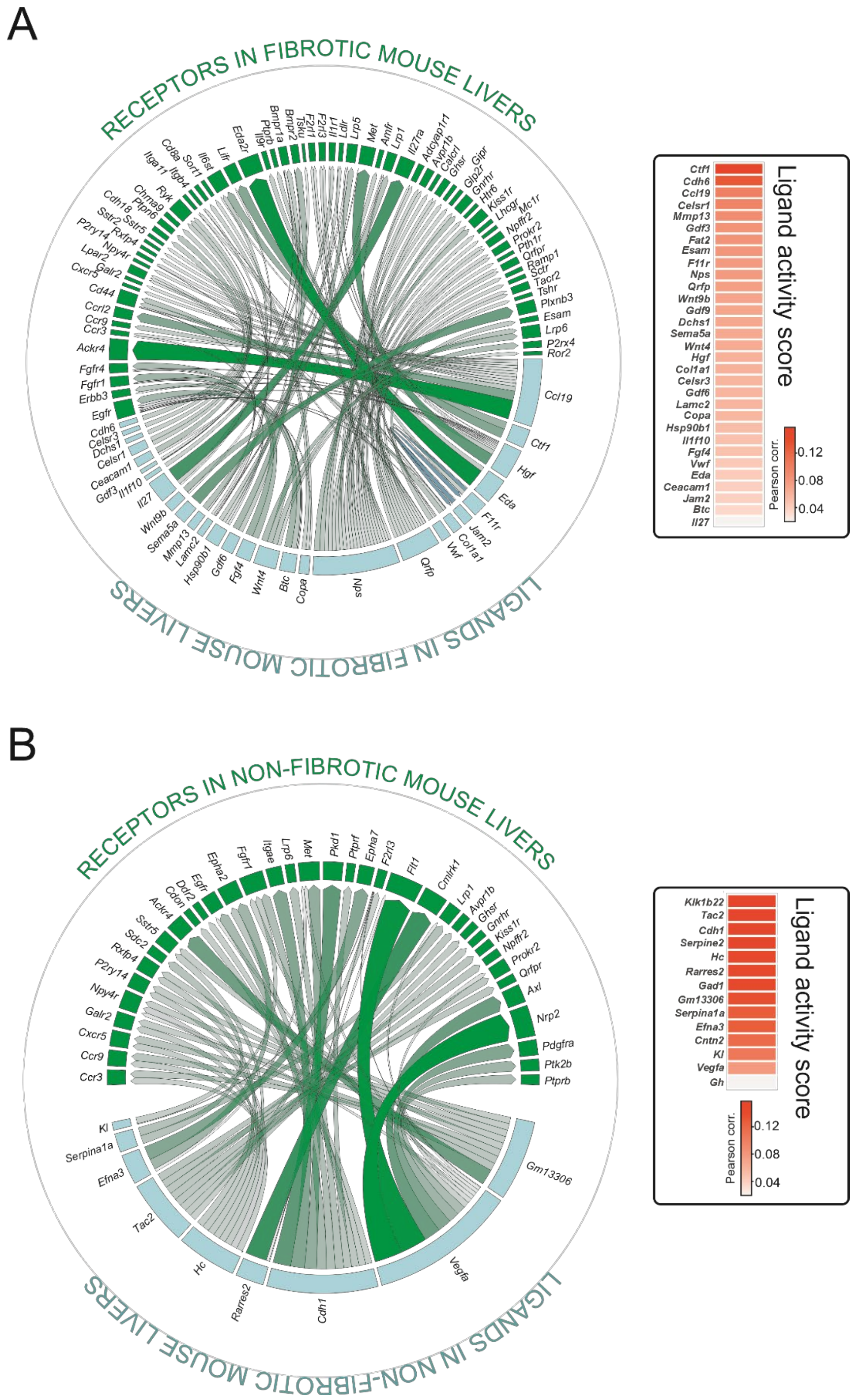

**Supplemental Figure 12: Ligand-receptor interactions in diet-driven MASH/fibrosis.** Gene expression in livers from chow- or "MASH diet"-fed male mice was assessed by RNA-seq and ligand-receptor (L-R) productive interactions were analyzed using the R package NicheNet adapted here for bulk tissue characterization. Relevant ligand-receptor pairs and associated ligand activity score were prioritized according to their predicted ability to account for the observed gene-expression changes. A) L-R interactions and ligand activity scores in livers from "MASH" diet-fed mice. B) L-R interactions and ligand activity scores in livers from chow diet-fed (control) mice.

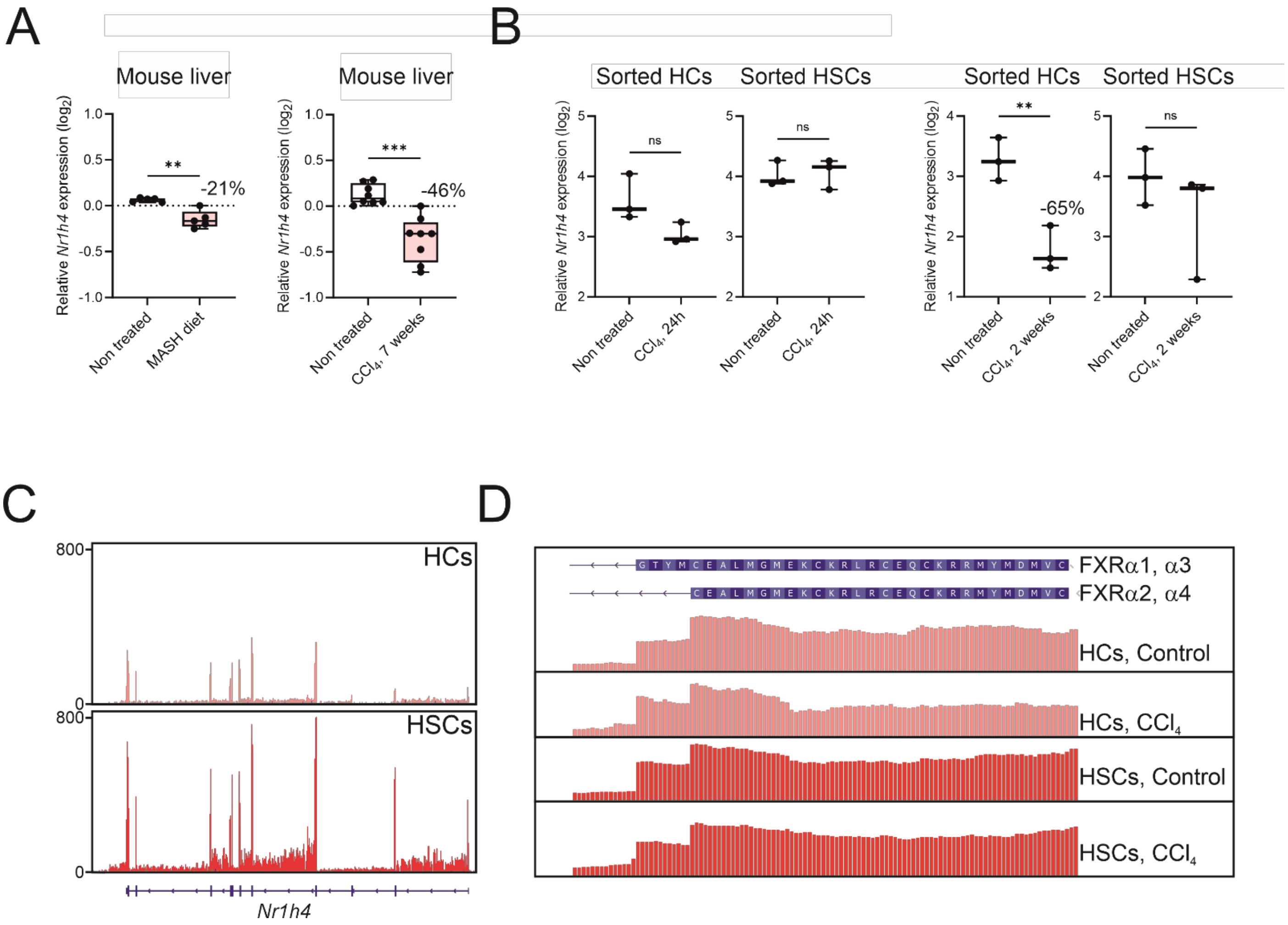

**Supplemental Figure 13: *Nr1h4* expression and splicing in mouse liver.** A) *Nr1h4* gene expression in mouse liver after MASH diet feeding or $CCl_4$ exposure. Gene expression was measured by RT-qPCR. Difference between groups were compared using an unpaired two-tailed t-test (p<0.01: **; p<0.005: ***). B) *Nr1h4* gene expression in mouse liver after a short $CCl_4$ exposure. Gene expression was measured by RT-qPCR. Difference between groups were compared using an unpaired two-tailed t-test (p<0.01: **). C) RNA-seq read coverage across the *Nr1h4* locus in HCs and HSCs isolated from unchallenged mouse livers. Reads were visualized using IGV. D) RNA-seq reads spanning the α1, α3/α2, α4 splice junction in unchallenged or $CCl_4$-exposed HCs and HSCs. Sequences are shown in the reverse orientation (read from right to left).

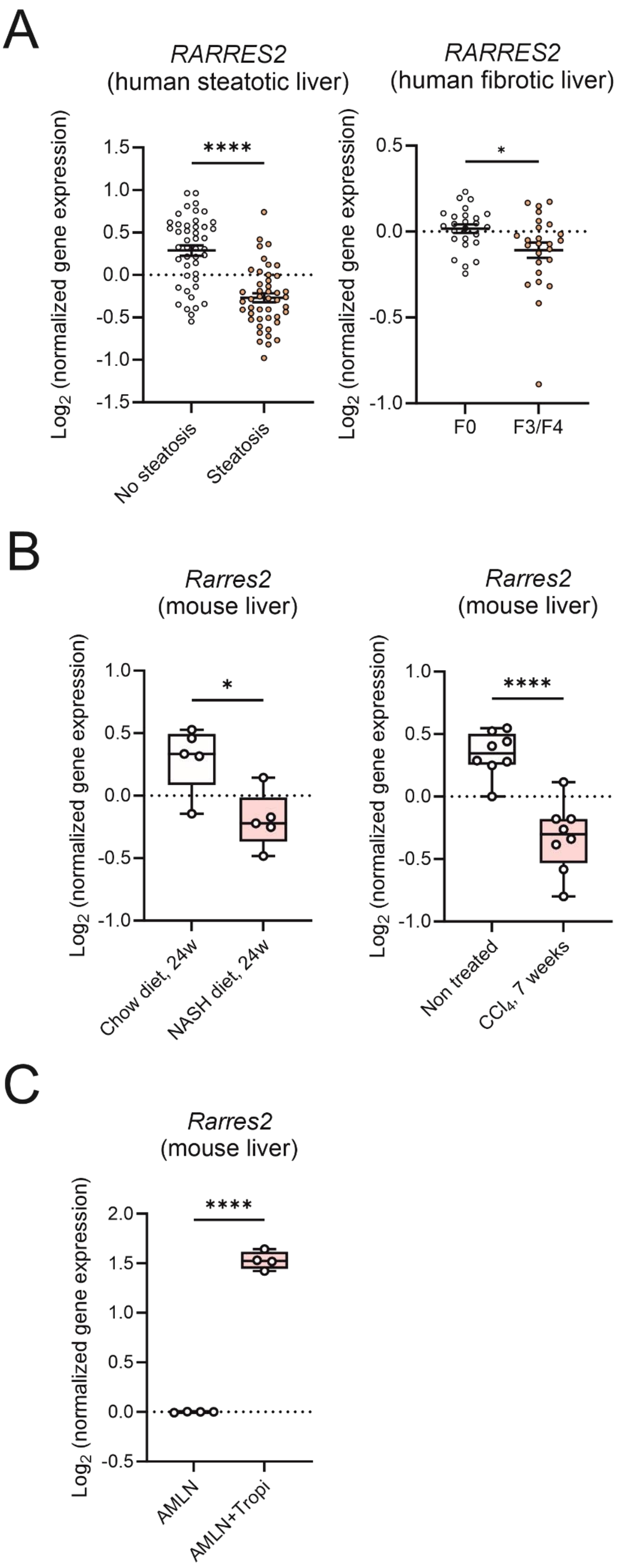

**Supplemental Figure 14: *Rarres2* expression in human and mouse liver.** A) *RARRES2* expression in human steatotic and fibrotic livers. Based on histological features, human liver biopsies were classified as non-steatotic or steatotic (grade > 1), and as non-fibrotic (F = 0) or fibrotic (Kleiner score F > 2). Transcriptomic profiles from propensity-matched biopsies (matched for age, sex, BMI, diabetic status, and statin use) were compared using DNA microarrays, and *RARRES2* expression values were extracted and analyzed using a two-tailed Welch's t-test (*p < 0.05; ****p < 0.001). B) *Rarres2* expression in mouse liver. Expression values for *Rarres2* were extracted from RNA-seq data obtained from control, MASH diet-fed, and $CCl_4$-exposed mouse livers. Differences between groups (n = 4–8) were analyzed using an unpaired two-tailed t-test (*p < 0.05; ****p < 0.001). C) *Rarres2* expression in livers from mice fed an Amylin (AMLN) diet for 26 weeks and treated or not with tropifexor (0.3 mpk) during the last 4 weeks. Difference among means (n=4) were compared using a unpaired two-tailed t-test (p<0.001: ****). RNA-seq data are from [28].

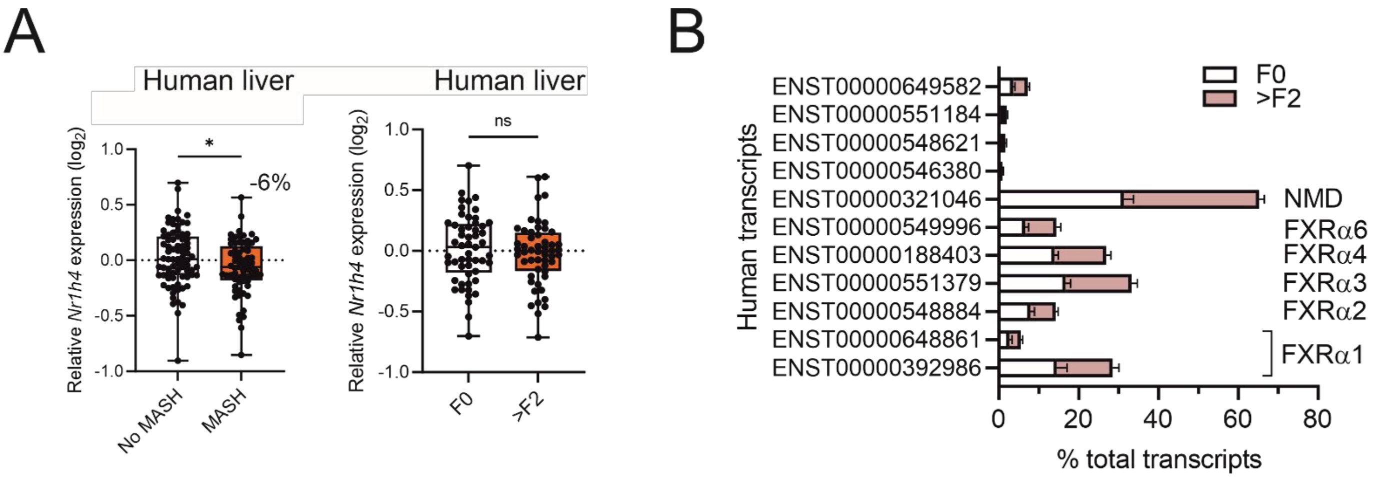

**Supplemental Figure 15: *NR1H4* expression and splicing in human livers.** A) *NR1H4* expression in human MASH and fibrotic livers. Based on histological features, human liver biopsies were classified as harboring MASH (steatosis grade >1, ballooning or lobular inflammation > 0) or not, and as non-fibrotic (F = 0) or fibrotic (Kleiner score F > 2). Transcriptomic profiles from propensity-matched biopsies (matched for age, sex, BMI, diabetic status, and statin use) were compared using DNA microarrays, and *NR1H4* expression values were extracted and analyzed using a two-tailed Welch's t-test (*p < 0.05). B) *NR1H4* isoforms in human histologically normal and fibrotic livers. RNA-seq data were processed to identify *NR1H4* isoforms.

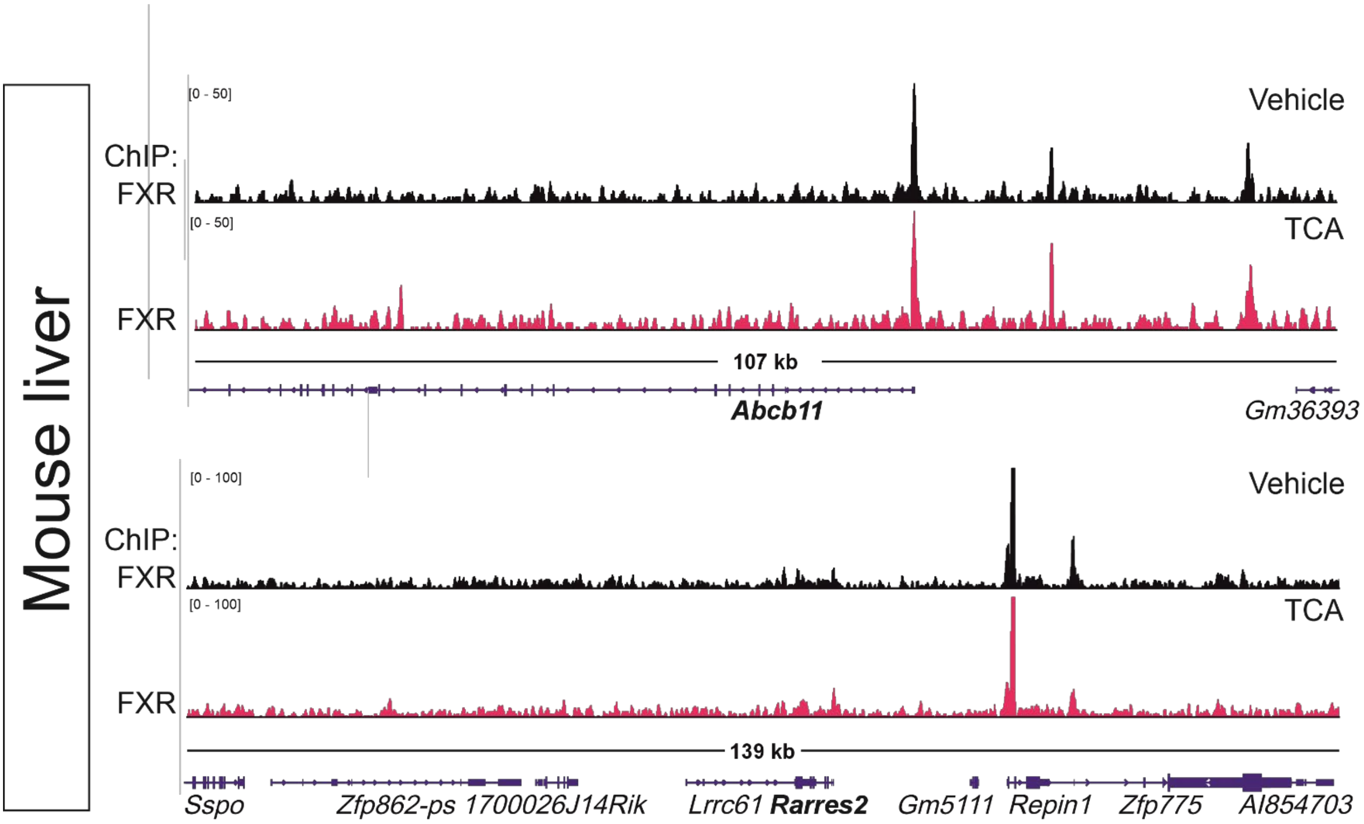

**Supplemental Figure 16: FXR ChIP-seq profile in mouse liver at the *Abcb11*/*Bsep* and *Rarres2* loci.** FXR ChIP-seq data from vehicle- or taurocholic (TCA)-exposed male mice [32] were visualized using IGV.

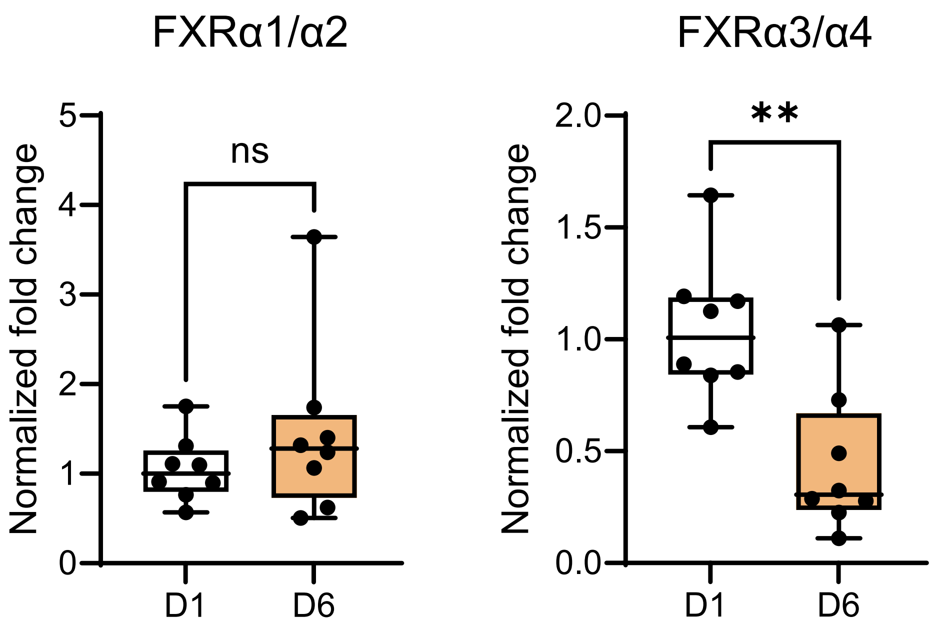

**Supplemental Figure 17: *Nr1h4* mRNA splice variants in PCLS.** FXR isoform content was assayed by RT-QPCR using primers specific for the α1/α2 and α3/α4 isoforms. Differences between groups (n = 8) were analyzed using an unpaired two-tailed t-test (*p < 0.05; ****p < 0.001).

A

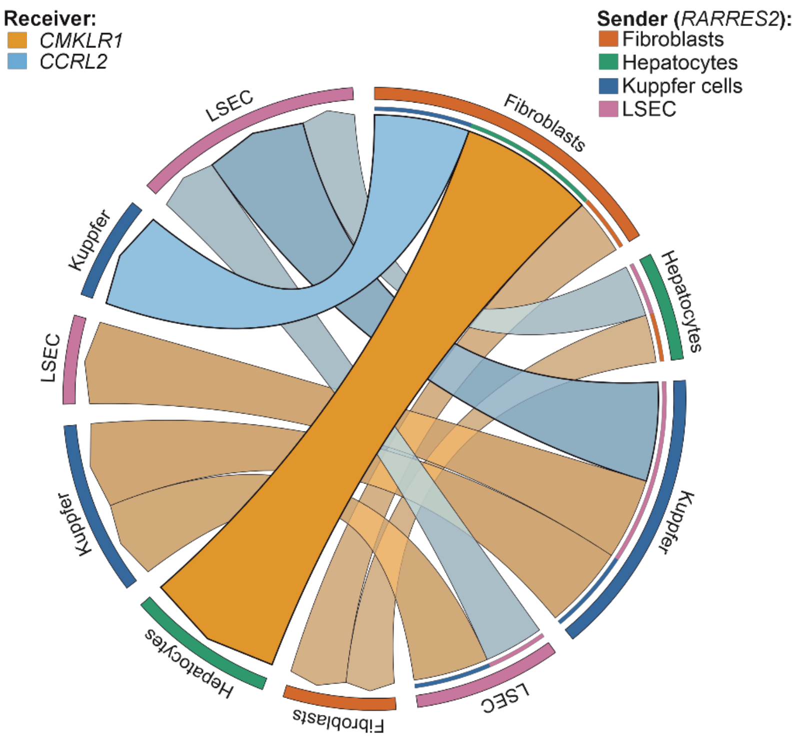

B

| Interaction | Source | Target | Spearman correlations of ligand and receptor zonation profiles |
|---|---|---|---|
| **Fibroblasts-Hepatocytes** | RARRES2 | CMKLR1 | 0,81 |
| **Kupffer-LSEC** | RARRES2 | CCRL2 | 0,60 |
| **Fibroblasts-Kupffer** | RARRES2 | CCRL2 | 0,57 |
| Kupffer-Kupffer | RARRES2 | CMKLR1 | 0,50 |
| Kupffer-LSEC | RARRES2 | CMKLR1 | 0,49 |
| LSEC-Kupffer | RARRES2 | CMKLR1 | 0,45 |
| LSEC-LSEC | RARRES2 | CCRL2 | 0,36 |
| Fibroblasts-Fibroblasts | RARRES2 | CMKLR1 | 0,33 |
| Hepatocytes-LSEC | RARRES2 | CCRL2 | 0,31 |
| Hepatocytes-Fibroblasts | RARRES2 | CMKLR1 | 0,29 |
| Hepatocytes-Fibroblasts | RARRES2 | CCRL2 | 0,25 |
| Fibroblasts-LSEC | RARRES2 | CMKLR1 | 0,24 |
| Hepatocytes-Kupffer | RARRES2 | CCRL2 | 0,24 |
| Hepatocytes-Kupffer | RARRES2 | CMKLR1 | 0,21 |
| Kupffer-Kupffer | RARRES2 | CCRL2 | 0,19 |
| Fibroblasts-Kupffer | RARRES2 | CMKLR1 | 0,17 |
| Hepatocytes-Hepatocytes | RARRES2 | CCRL2 | 0,05 |
| LSEC-LSEC | RARRES2 | CMKLR1 | -0,07 |
| Hepatocytes-LSEC | RARRES2 | CMKLR1 | -0,10 |
| Kupffer-Hepatocytes | RARRES2 | CCRL2 | -0,14 |
| Kupffer-Hepatocytes | RARRES2 | CMKLR1 | -0,17 |
| Hepatocytes-Hepatocytes | RARRES2 | CMKLR1 | -0,21 |
| LSEC-Hepatocytes | RARRES2 | CCRL2 | -0,24 |
| Kupffer-Fibroblasts | RARRES2 | CCRL2 | -0,25 |
| LSEC-Fibroblasts | RARRES2 | CCRL2 | -0,25 |
| Kupffer-Fibroblasts | RARRES2 | CMKLR1 | -0,33 |
| LSEC-Hepatocytes | RARRES2 | CMKLR1 | -0,33 |
| Fibroblasts-Hepatocytes | RARRES2 | CCRL2 | -0,36 |
| Fibroblasts-Fibroblasts | RARRES2 | CCRL2 | -0,41 |
| LSEC-Kupffer | RARRES2 | CCRL2 | -0,48 |
| LSEC-Fibroblasts | RARRES2 | CMKLR1 | -0,55 |

**Supplemental Figure 18: Spatial concordance between *RARRES2* and its cognate receptors across zonated human liver cell types.** Spearman correlation coefficients between the zonation profile of *RARRES2* expression in sender cell types and those of *CMKLR1* or *CCRL2* expression in receiver cell types were derived from the dataset of Yakubovsky et al., 2026 [33]. Interactions showing positive spatial concordance above the selected threshold ($\rho > 0.25$) are displayed as a Circos plot (A). In the Circos plot, each chord links a sender cell type expressing *RARRES2* to a receiver cell type expressing *CMKLR1* or *CCRL2*; chord width is proportional to the Spearman correlation coefficient between ligand and receptor zonation profiles, and chord color denotes receptor identity. Color intensity also scales with the correlation coefficient. Positive correlations indicate concordant zonation patterns compatible with zone-specific intercellular communication. The complete set of pairwise correlations is provided in B).

**Supplemental table 1: Differential gene expression in wild type and liver (HC)-specific FXR KO mice.** RNA-seq data from livers from male mice of the 2 indicated genetic backgrounds were analyzed as described in the Materials and Methods section to generate a list of significantly differentially expressed genes (Sheet 1). RPKM values for each liver cell types are indicated in sheet 2 (taken from Zummo et al., 2023 [6]).

**Supplemental table 2: Differential gene expression in livers from male mice treated with OCA or tropifexor.** RNA-seq data from livers from wild-type male mice were analyzed as described in the Materials and Methods section to generate a list of significantly differentially expressed genes following short term (3 days) treatment with either compound.

**Supplemental table 3: Differential gene expression in purified hepatocytes or hepatic stellate cells from male mice treated with tropifexor.** RNA-seq data from HCs or HSCs isolated from wild-type male mice treated with tropifexor for 3 days were analyzed as described in the Materials and Methods section to generate a list of significantly differentially expressed genes in each cell type.

**Supplemental table 4: Cell-specific expression of ligand-receptor pairs in liver cell types in response to tropifexor.** RNA-seq data from tropifexor-exposed HCs or HSCs were extracted from Supplemental table 3. Differential gene expression in response to treatment, as well as cell-specific expression levels is indicated for each ligand or receptor.

**Supplemental table 5: Differential gene expression in response to tropifexor in precision-cut liver slices, livers from wild-type mice, and livers from AMLN diet-fed mice.** RNA-seq data from tropifexor-exposed PCLS, livers from wild-type mice, or livers from AMLN diet-fed mice were analyzed as described in the Materials and Methods section to generate a list of significantly differentially expressed genes in each experimental system. Data from AMLN diet-fed mice were obtained from[28].
